# K29-Linked Ubiquitin Signaling Regulates Proteotoxic Stress Response and Cell Cycle

**DOI:** 10.1101/2020.10.15.341719

**Authors:** Yuanyuan Yu, Qingyun Zheng, Satchal K. Erramilli, Man Pan, Seongjin Park, Yuan Xie, Jingxian Li, Jingyi Fei, Anthony A. Kossiakoff, Lei Liu, Minglei Zhao

## Abstract

Protein ubiquitination shows remarkable topological and functional diversity through the polymerization of ubiquitin via different linkages. Deciphering the cellular ubiquitin code is of central importance to understand the physiology of the cell. Among the eight possible linkages, K29-linked polyubiquitin is a relatively abundant type of polyubiquitin in both human and yeast cells. However, our understanding of its function is rather limited due to the lack of specific binders as tools to detect K29-linked polyubiquitin. In this study, we screened and characterized a synthetic antigen-binding fragment, termed sAB-K29, that can specifically recognize K29-linked polyubiquitin using chemically synthesized K29-linked diubiquitin. We further determined the crystal structure of this fragment bound to the K29-linked diubiquitin, which revealed the molecular basis of specificity. Using sAB-K29 as a tool, we uncovered that K29-linked ubiquitination is involved in different kinds of cellular proteotoxic stress response as well as cell cycle regulation. In particular, we showed that K29-linked ubiquitination is enriched in the midbody and downregulation of the K29-linked ubiquitination signal arrests cells in G1/S phase.

## INTRODUCTION

Protein ubiquitination is a dynamic posttranslational modification involved in nearly all aspects of eukaryotic cells^1–3^. Ubiquitin, a 76-amino acid protein, can be covalently attached to its substrate through an isopeptide bond between its C-terminal carboxyl group and the side-chain amino group of a lysine residue on the substrate, initiating the process of ubiquitination. The substrate protein can be modified by either a single ubiquitin, called monoubiquitination, or a chain of ubiquitin molecules, termed polyubiquitination. To form a ubiquitin chain, the second and subsequent ubiquitin molecules need to be conjugated to one of the seven lysine residues (K6, K11, K27, K29, K33, K48, and K63) or the N-terminal amino group of the first methionine residue (M1) on the preceding ubiquitin molecule (**Fig. 1a**). Hence, a total of eight linkages between any of two covalently linked ubiquitin molecules exist in vivo, which makes polyubiquitination an extremely diverse posttranslational modification^1^. Recently, branched and heterotypic ubiquitin chains were also discovered in eukaryotic cells, further expanding the complexity of ubiquitination^4–6^. Ubiquitin chains with different linkages have different topologies and structures, creating a multitude of distinct signals, known as the ubiquitin code.

**Fig. 1.**
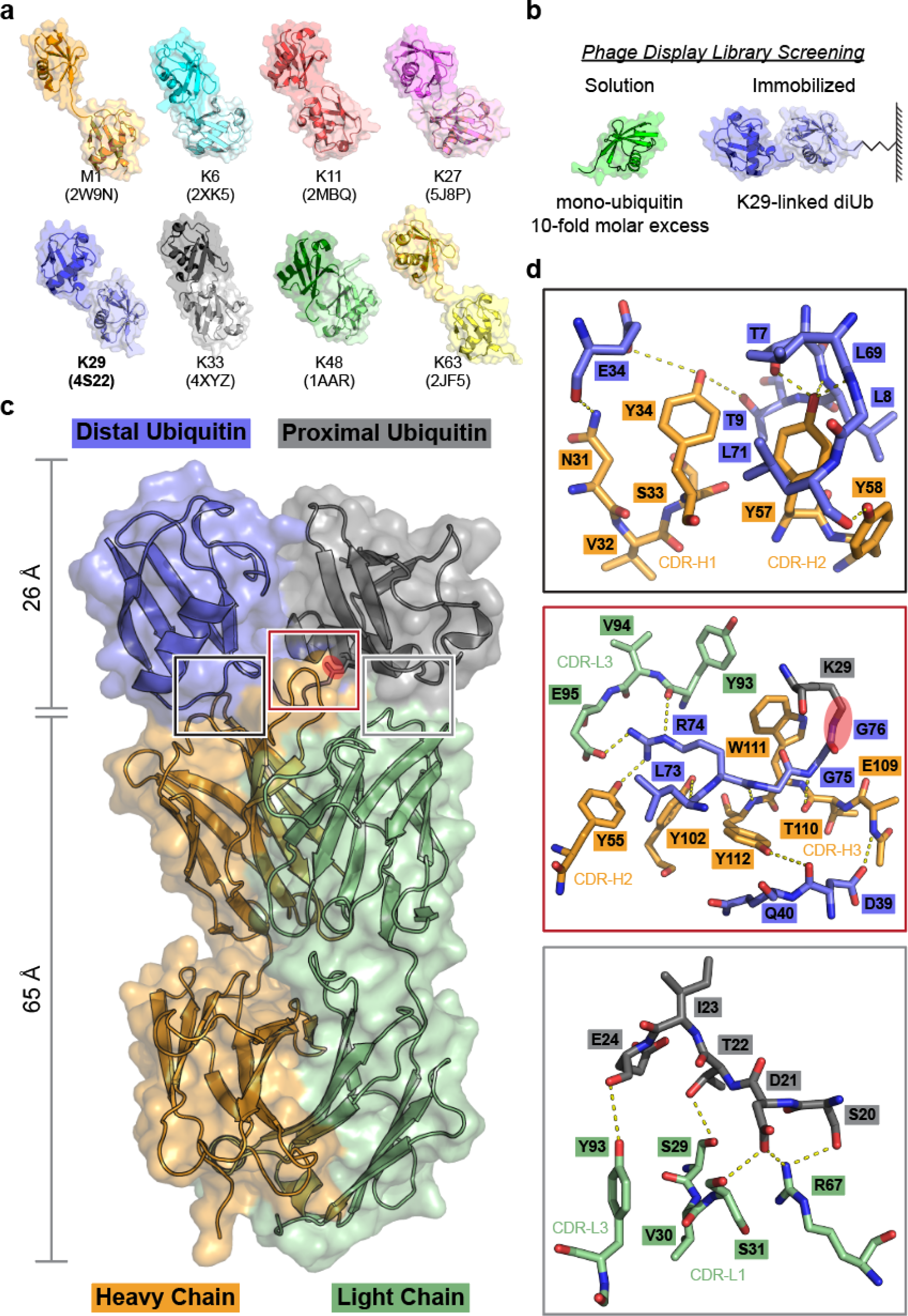
Screening for a K29-linked ubiquitin chain-specific binder and the complex structure of sAB-K29 and K29-linked diubiquitin. **a**, Crystal structures of diubiquitin molecules with eight possible linkages. The K29-linked ubiquitin chain was the focus of this study. **b**, Screening of a phage display library of engineered Fabs using immobilized synthetic K29-linked diubiquitin. Monoubiquitin at a 10-fold molar excess was used for negative selection. **c**, Crystal structure of sAB-K29 in complex with K29-linked diubiquitin determined at a 2.9 Å resolution (**Extended Data Table 1**). Color code: orange, heavy chain of sAB-K29; green, light chain of sAB-K29; blue, distal ubiquitin; gray, proximal ubiquitin. The isopeptide bond between the C-terminus of the distal ubiquitin molecule and K29 of the proximal ubiquitin molecule is highlighted with a red oval. **d**, Zoomed-in views of the three binding interfaces between sAB-K29 and the diubiquitin. Top, binding interface between the heavy chain of sAB-K29 and the distal ubiquitin molecule; middle, binding interface between both the heavy and light chains of sAB-K29 and the linker region of the diubiquitin molecule; bottom, binding interface between the light chain of sAB-K29 and the proximal ubiquitin molecule. Potential hydrogen bonds are shown as dotted lines. The isopeptide bond in the middle panel is highlighted with a red oval.

Deciphering cellular ubiquitin code is of central interest in ubiquitin studies^7–9^. For instance, K48-linked polyubiquitin chains adopt a closed conformation and target their substrate proteins to the proteasome for degradation^10–12^. In contrast, K63-linked polyubiquitin chains adopt an open conformation and regulate proteasome-independent signaling pathways such as DNA repair, inflammatory signaling, and endocytosis^13^. Other types of polyubiquitin chains, regarded as atypical, have also been reported to adopt distinct structures and perform specific cellular functions^14–16^. For example, M1-linked ubiquitin precipitates in immune response^17–19^. K6-linked ubiquitin regulates Mitophagy^20–23^. K11-linked ubiquitin is involved in cell cycle regulation and protein degradation^23, 24^. K27-linked ubiquitin regulates autoimmunity, innate immune response and tumorigenesis^25–27^. K33-linked ubiquitin mediates protein trafficking and signal transduction of cell surface receptors^28, 29^. Remarkably, the function of K29-linked ubiquitin is much less understood compared to the other topologies, although it has been reported involving in protein degradation and the regulation of a viral infection^30, 31^. It is worth noting that, according to a recent quantitative study in eukaryotic cells^32^, the abundance of K29-linked ubiquitin is the highest among the atypical linkage types, close to K63-linked ubiquitin, and follow only K48-linked ubiquitin. The limited knowledge of K29-linked ubiquitin is largely due to a lack of linkage-specific detection tools. Previously, recombinant TRABID NZF1-Halo fusions coupled with vOTU treatment have been used to study K29-linked polyubiquitin chains^33^. However, the procedure involved using vOTU, a deubiquitinating enzyme (DUB), to remove other linkages and therefore had limited applications. All the other homotypic ubiquitin chains have linkage-specific detection tools^34^. Newton et al. developed two linkage-specific antibodies that specifically recognize K48- and K63-linked polyubiquitin chains and used them to reveal that polyubiquitin editing is a general mechanism to attenuate innate immune signaling^35^. Matsumoto et al. engineered a K11 linkage-specific antibody and used it to demonstrate that K11-linked polyubiquitin are critical regulators for the degradation of mitotic proteins^23^. Matsumoto et al. also engineered a linear (M1) linkage-specific antibody^36^, which was used to uncover the role of linear chains in immune responses. Michel et al. characterized linkage-specific “affimer” reagents that recognize K6- and K33-linked polyubiquitin chains and used the K6 affimer to show that HUWE1 is the main E3 ligase for the K6-linked polyubiquitination of mitofusin-2^34^. For a heterotypic polyubiquitin chain, Yau et al. engineered a bispecific antibody that recognizes K11/K48-linked chains and used it to show that mitotic regulators, misfolded nascent polypeptides, and pathological Huntingtin variants are endogenous substrates of K11/K48-linked polyubiquitination^37^.

In this study, we developed and characterized a synthetic antigen-binding fragment (sAB) that can specifically recognize K29-linked ubiquitin chains at nanomolar concentrations by screening a phage display library. We determined the crystal structure of the sAB in complex with K29-linked diubiquitin (diUb) and elucidated the specific interactions between the two components around the isopeptide bond. Through a series of experiments, including pull-down assays, mass spectrometry and immunofluorescent imaging, we revealed the interaction landscape of K29-linked ubiquitination which showed that it is widely involved in protein homeostasis, stress response, and cell cycle regulation. Interestingly, we observed that K29-linked ubiquitination is enriched in puncta under several proteotoxic cellular stresses including unfolded protein response, oxidative stress and heat shock response. Moreover, we found that K29-linked ubiquitination was particularly enriched in the midbody at the telophase of mitosis. The knockdown of K29-linked ubiquitination by a specific DUB led to arrest of the cell cycle at G1/S phase. In summary, we have established that K29-linked ubiquitin chains are important players in stress response and cell cycle signaling pathways.

## RESULTS

### Selection of K29-linked polyubiquitin-specific sAB (sAB-K29) from a phage display library

sABs have been successfully developed as linkage-specific binders for M1-, K11-, K48-, and K63-linked ubiquitin, as mentioned above. They are functionally and structurally similar to antigen-binding fragments (Fabs) and can be selected from a synthetic phage display library ^38^. To obtain a K29-linked ubiquitin-specific binder, biotinylated K29-linked diUb without any other linkages is required. Therefore we chose to chemically synthesize biotinylated K29-linked diUb following previously developed methods ^39^. A polyethylene glycol (PEG) linker was introduced between the diUb and biotin moieties. The synthetic route is briefly summarized **(Extended Data Fig.1a**). We conducted reverse-phase high-performance liquid chromatography (RP-HPLC) followed by liquid chromatography-mass spectrometry (LC-MS) to verify the mass of the product (**Extended Data Fig.1b**). Next, we refolded the product using a standard dialysis method and collected the circular dichroism (CD) spectrum of the final product (**Extended Data Fig.1c**). The CD spectrum of the synthesized K29-linked diUb showed an ellipticity near identical to that of the ubiquitin monomer (monoUb), suggesting correct folding of the diUb ^40^.

We subsequently generated a K29-linked diUb-specific sAB from a phage display library, known as Library E, based on a humanized antibody Fab scaffold ^41^. This in vitro technique allows for exquisite control over sAB selection conditions ^42^. We took advantage of this methodology by using an excess of monoUb in solution (**Fig. 1b**) to generate a linkage-specific binder, which we termed sAB-K29 and characterized further as described below.

### Structural characterization of sAB-K29 in complex with K29-linked diUb

To enable crystal screening of K29-linked diUb and sAB complex at a high concentration, a large quantity of K29-linked diUb was required. We therefore used the enzymatic preparation method ^33^. Although the linkage of the diUb is not as pure as the chemical approach, only K29-linked diUb could form a complex with sAB-K29. Specifically, ubiquitin was mixed with UBA1, UBE2L3, and UBE3C for polyubiquitin chain formation. The resulting polyubiquitin was a mixture of mainly K48- and K29-linked chains. vOTU, a DUB that does not cleave K29-linked chains, was introduced to the mixture to remove the K48-linked chains. Finally, K29-linked diUb was separated from monoUb and polyubiquitin using anion exchange chromatography (**Extended Data Fig. 2a**). To obtain the complex of sAB-K29 and K29-linked diUb, we mixed purified K29-linked diUb with sAB-K29 at a molar ratio of 1.5:1 and ran the mixture through gel filtration (**Extended Data Fig. 2b and c**). The stable complex was subjected to crystallization screening (see methods for details). Diffraction-quality crystals appeared from a reservoir solution containing 0.1 M phosphate-citrate and 40% v/v PEG 300 (**Extended Data Fig. 2d**). Diffraction data to 2.9 Å resolution were collected at the Advanced Photon Source at Argonne National Lab (**Extended Data Table 1**).

**Fig. 2.**
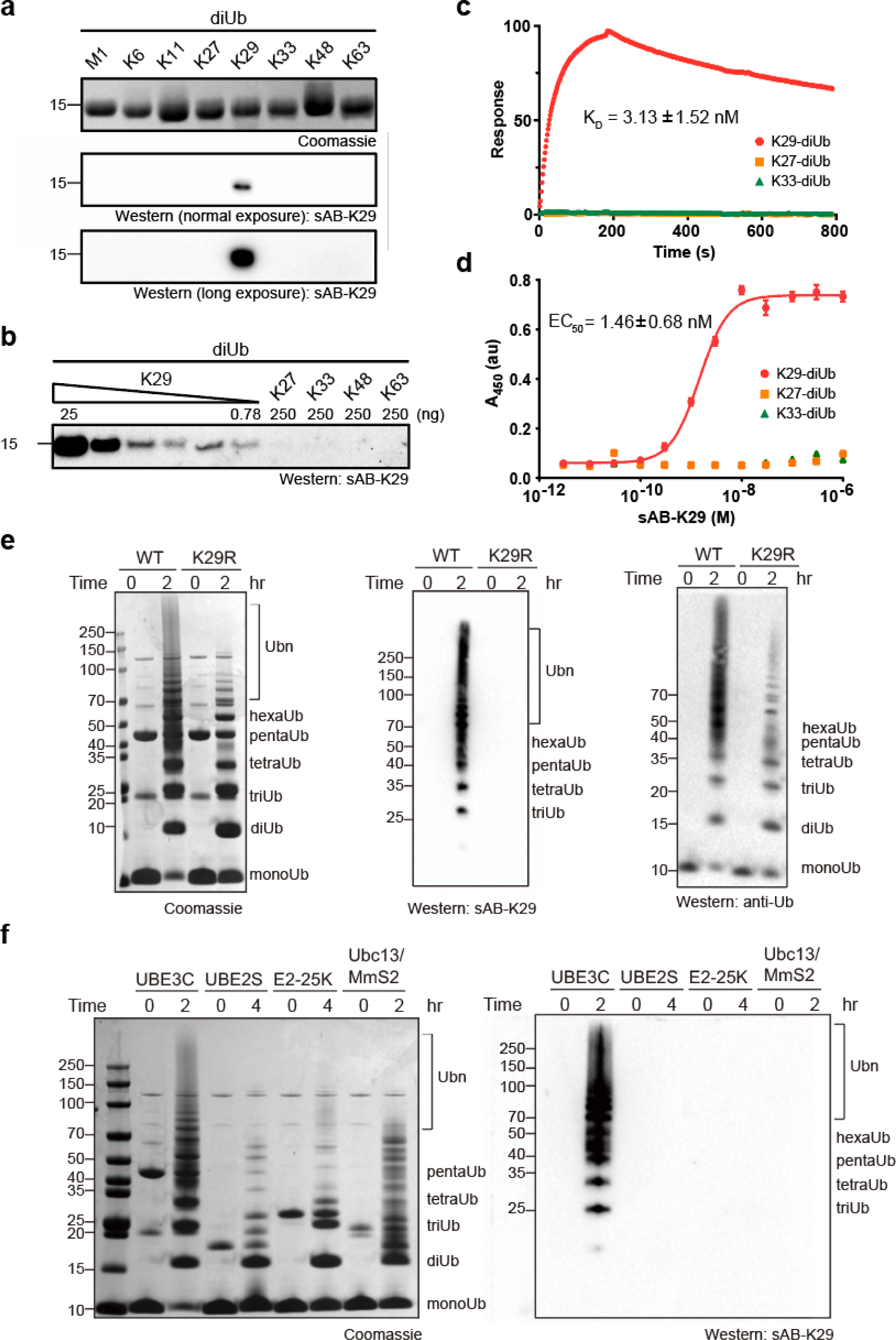
In vitro characterization of sAB-K29. **a**, The specificity of sAB-K29. Diubiquitin molecules (∼500 ng) with eight types of linkages were loaded on an SDS-PAGE gel. Western blotting was performed using sAB-K29 as the primary binder and peroxidase-conjugated goat anti-human IgG (F(ab)2 fragment-specific, Jackson ImmunoResearch) as the secondary antibody under normal or long exposure. **b**, The sensitivity and crossreactivity of sAB-K29. sAB-K29 could detect K29-linked diUb at the level of 0.78 ng in a Western blot and had no crossreactivity with K27/K33/K48/K63-linked diUb at the level of 250 ng. **c**, Surface plasma resonance (SPR) experiments using immobilized K29/K27/K33-linked diUb and purified sAB-K29. The measured dissociation constant (K_D_) between K29-linked diUb and sAB-K29 was 3.13 ± 1.52 nM. No binding between sAB-K29 and K27/K33-linked diUb was detected. **d**, ELISAs using immobilized K29/K27/K33-linked diUb and a series of 12 serially diluted purified sAB-K29 solutions. The measured EC_50_ was 1.46 ± 0.68 nM for K29-linked diUb. No significant absorption was detected for K27/K33-linked diUb. **e**, sAB-K29 could be used to detect K29-linked polyubiquitin chains assembled in vitro by Western blotting. K29-linked polyubiquitin chains were assembled by an E1-E2-E3 mixture containing UBA1, UBE2L3, and UBE3C. Parallel Western blotting using anti-ubiquitin antibody was performed for comparison. The ubiquitin mutant K29R was used in the control experiment. **f**, sAB-K29 could specifically detect K29-linked polyubiquitin chains in vitro. Polyubiquitin chains were assembled by four ubiquitination systems: UBE3C (K48- and K29-linked chains), UBE2S (K11-linked chains), E2-25K (K48-linked chains), and UBC13/Mms2 (K63-linked chains).

The complex structure shows a 1:1 stoichiometry of sAB-K29 and K29-linked diUb (**Fig. 1c**). There are three binding interfaces between the complementarity-determining regions (CDRs) of sAB-K29 and diUb (**Fig. 1c**). The left interface (black box) involves the heavy chain of sAB-K29 (CDR-H1 and H2) and the distal ubiquitin molecule. The right interface (gray box) involves the light chain of sAB-K29 (CDR-L1 and L3) and the proximal ubiquitin molecule. The middle interface (red box) involves both chains (CDR-H2, H3 and L3) and the linker between the two ubiquitin molecules. Hydrogen bonding networks and van der Waals interactions mainly contributed by tyrosine and serine residues dominate the interfaces (**Fig. 2d**). Each of the three interfaces recognizes an essential piece of the diUb, namely, the proximal ubiquitin, distal ubiquitin, and linker. Together, they form the basis for specific recognition. Notably, the patch containing I44 on both ubiquitin molecules is not involved in the binding interface. Only the patch containing I36 of the distal ubiquitin molecule is involved in the interaction with the heavy chain of sAB-K29 (**Extended Data Fig. 2e**). Compared to the crystal structure of K29-linked diUb (PDB accession code: 4S22), the diUb in this complex adopts a more compact conformation, with the proximal ubiquitin molecule rotated approximately 60 degrees compared to that in the other structure (**Extended Data Fig. 2f**). Compared to the crystal structure of K29-linked diUb in complex with the NZF1 domain of TRABID (PDB accession code: 4S1Z), the diUb in 4S1Z is in a compact conformation similar to our structure, but the proximal ubiquitin is rotated about 90 degree. The NZF1 domain of TRABID binds at a different place, primarily recognizing the I44 patch of the distal ubiquitin (**Extended Data Fig. 2g**). We also superimposed K33-linked diUb (PDB accession code: 4XYZ) onto the diUb in our complex. The proximal ubiquitin of K33-linked diUb adopts a different orientation, which would not allow sAB-K29 to bind (**Extended Data Fig. 2h**).

### In vitro characterization of sAB-K29

We first performed single-point competitive phage enzyme-linked immunosorbent assay (ELISA) using M13 phage-displayed sAB-K29. The signal decreased when K29-linked diUb, but not K27- or K33-linked diUb, was used as the competitor in solution, suggesting its specificity towards K29-linked diUb (**Extended Data Fig. 4a**). Next, we tested the specificity of sAB-K29 against diUb of all linkage types (100 ng) by Western blotting. sAB-K29 recognized only K29-linked diUb and showed no cross-reactivity with other linkage types even after long exposure (**Fig. 2a**). We further performed Western blotting against serially diluted K29-linked diUb from 25 ng to 0.78 ng. The results suggested that sAB-K29 is at least 320-fold more sensitive for K29-linked diUb than the other diUbs tested (**Fig. 2b**). We measured the binding affinity of sAB-K29 and K29-linked diUb using surface plasmon resonance (SPR) and ELISA. By SPR, the dissociation constant (K_D_) was measured to be 3.13 ± 1.52 nM, which was in agreement with the EC50 of 1.46 ± 0.68 nM measured by ELISA.

Next, we tested whether sAB-K29 can be used to detect K29-linked polyubiquitin chains. We mixed ubiquitin with UBA1, UBE2L3, and UBE3C and monitored assembly of the polyubiquitin chain using Western blotting. UBE3C is known to produce a mixture of mainly K48- and K29-linked polyubiquitin chains ^43^. K29R mutant ubiquitin, which cannot form K29-linked polyubiquitin chains, was included as a control. The results of Western blotting showed that sAB-K29 specifically recognized K29-linked polyubiquitin chains (**Fig. 2f**, left), although the K29R mutant did not prevent the assembly of other linkages of polyubiquitin chains, as detected with a commercial anti-ubiquitin antibody and in a Coomassie blue stained gel (**Fig. 2e**). A single-point mutation in the heavy chain of sAB-K29, Y112A, could not recognize the K29-linked polyubiquitin chain under normal conditions, as shown by Western blotting (**Extended Data Fig. 3c).** We further assembled K11-, K48- and K63-linked polyubiquitin chains using UBE2S, E2-25K, and Ubc13/Mms2, respectively, and monitored the reactions using SDS-PAGE and Western blotting. Again, sAB-K29 could detect only the K29-linked polyubiquitin chains produced by the UBE3C system, not polyubiquitin chains with other types of linkages (**Fig. 2f, Extended Data Fig. 3d**). Together, these results suggest that sAB-K29 can be used as a tool to specifically detect K29-linked polyubiquitin chains in vitro.

**Fig. 3.**
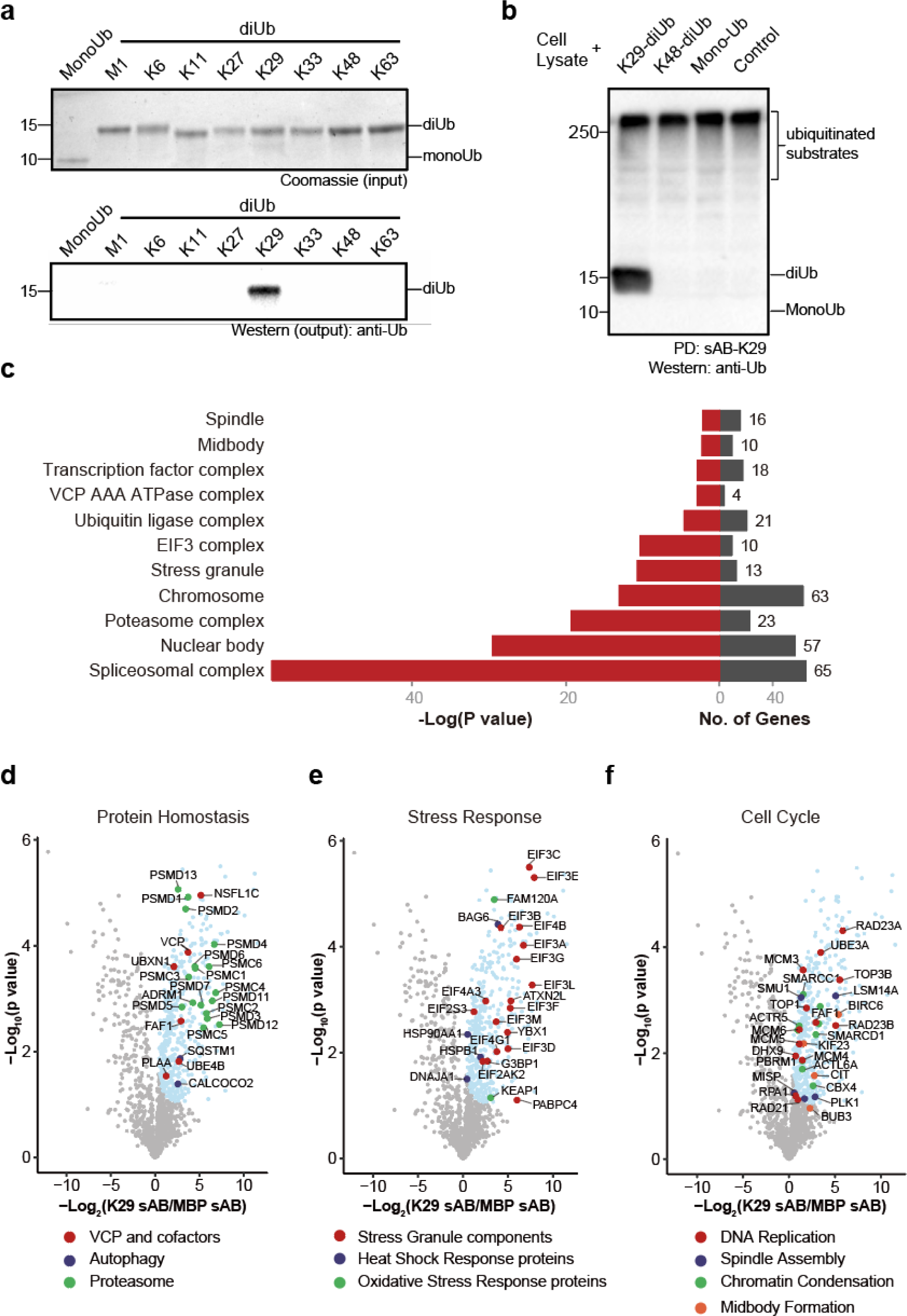
Pull-down and proteomic analyses of HeLa cells using sAB-K29. **a**, sAB-K29 could be used to specifically pull down K29-linked diUb in vitro. Biotinylated sAB-K29 was used to pull down mono- and diubiquitin with eight types of linkages. Anti-ubiquitin antibody was used in Western blotting of the output. **b**, sAB-K29 could be used to specifically pull down K29-linked diUb and K29-linked polyubiquitin chains from cell lysates. Lysates of HeLa cells were mixed with K29- and K48-linked polyubiquitin and monoubiquitin. Biotinylated sAB-K29 was used for the pull-down experiment. Anti-ubiquitin antibody was used for Western blotting of the output. **c**, GO analysis (cellular components) of enriched proteins from pull-down experiments of HeLa cells using sAB-K29. The enriched proteins were identified using label-free quantitative mass spectrometry. Three biological and two technical replicates were performed. Significant hits (compared to pull-down using sAB-MBP, FDR < 0.05) were subjected to GO analysis. **d-f**, Volcano plots of the quantitative mass spectrometry results. Significantly enriched hits (FDR < 0.05) are colored cyan, with some well-documented proteins involved in protein homeostasis (D), the stress response (E), and the cell cycle (F) highlighted and labeled.

**Fig. 4.**
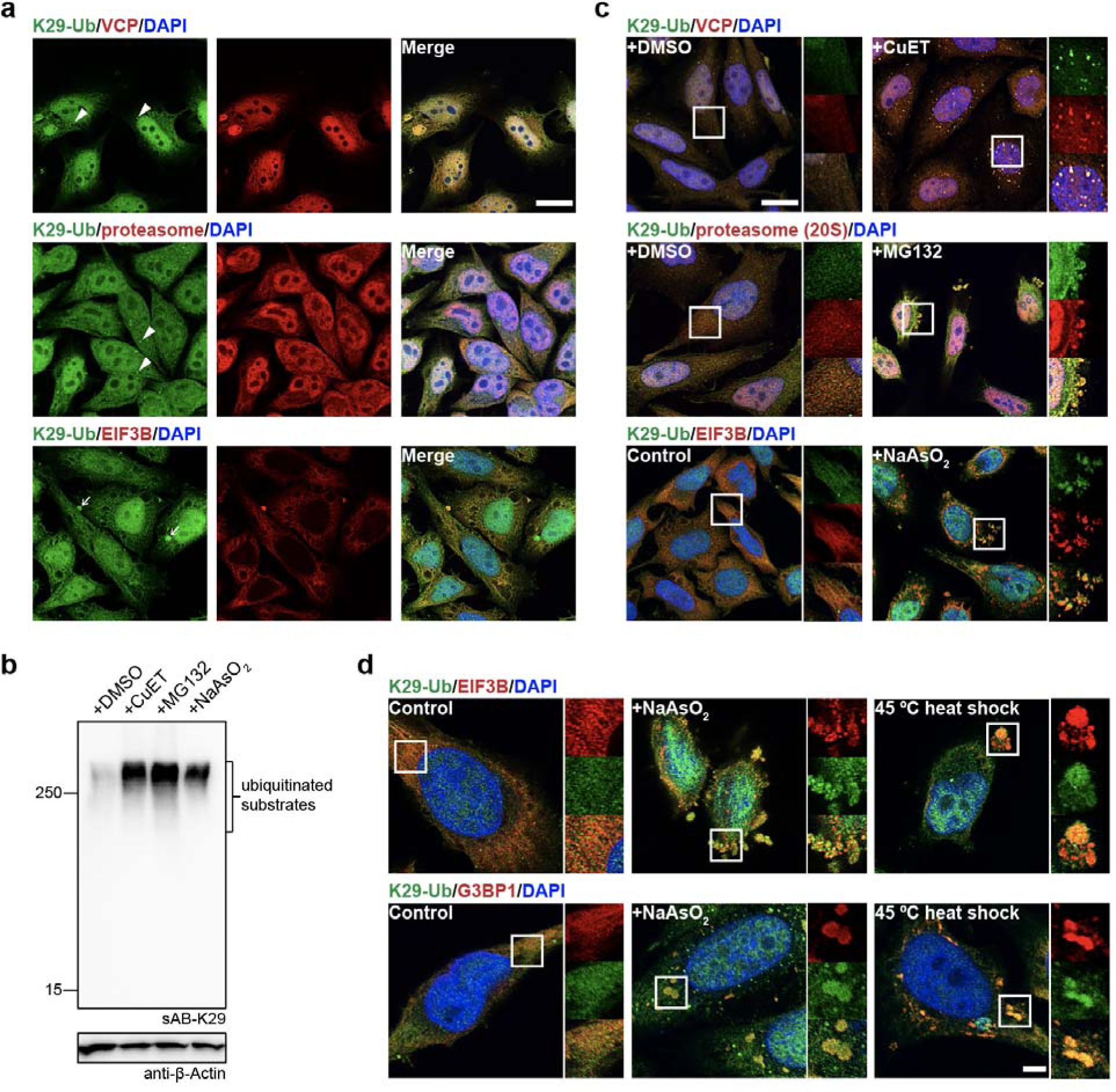
K29-linked polyubiquitination is involved in protein homeostasis and the stress response. **a**, sAB-K29 was used for the immunofluorescent staining of untreated HeLa cells. Costaining for K29-Ub with VCP, the proteasome (20S), and EIF3B is shown. Arrowheads point to bright puncta stained by sAB-K29. Arrows point to liquid droplets stained by sAB-K29. **b**, K29-linked polyubiquitination was enhanced after HeLa cells were treated with CuET (an inhibitor of the VCP cofactor Npl4), MG132 (a proteasome inhibitor), or sodium arsenite (a stress response inducer). HeLa cells were treated with 1 μM CuET for 4 h, 20 μM MG132 for 4 h, or 500 μM NaAsO_2_ for 1 h, followed by Western blot analysis using sAB-K29 as the primary binder. **c**, Immunofluorescent staining of HeLa cells treated with 1 μM CuET for 4 h, 100 μM MG132 for 4 h, or 500 μM sodium arsenite for 1 h. Costaining for K29-Ub with VCP, the proteasome (20S), and EIF3B is shown. **d**, K29-linked polyubiquitination was enriched in stress granules. HeLa cells were treated with 500 μM sodium arsenite for 1 h or heat shocked at 45 °C for 30 minutes, followed by immunofluorescent staining. Costaining for K29-Ub with EIF3B and G3BP1 is shown. Scale bars in panels **a** and **c**: 20 μm, panel **d**: 5 μm.

### K29-linked ubiquitination is involved in protein homeostasis, RNA processing, the stress response, and cell cycle regulation

To further investigate the function of K29-linked ubiquitination in cells, we introduced a biotin tag at the C-terminus of the heavy chain of sAB-K29 using sortase-catalyzed ligation ^44, 45^. A polyethylene glycol (PEG) linker between the biotin moiety and sAB-K29 was introduced to increase the flexibility. Using biotinylated sAB-K29, we first performed pull-down experiments against monoUb and diUb of all linkage types. The results showed that K29-linked diUb was specifically pulled down (**Fig. 3a**). Next, we tested whether the pull-down assay would work in a crowded environment by mixing HeLa cell lysate with K29-linked diUb, K48-linked diUb, and monoUb and performed Western blotting using an anti-ubiquitin antibody. Only K29-linked diUb was pulled down with high-molecular-weight endogenous species (**Fig. 3b**), suggesting that biotinylated sAB-K29 can be used as a specific tool for cellular pull-down assays.

To elucidate the function of K29-linked ubiquitination, we conjugated biotinylated sAB-K29 to magnetic streptavidin beads and used the beads to capture K29-linked ubiquitinated proteins from HeLa cell lysates. The enriched proteins were eluted with 8 M urea and subjected to label-free mass spectrometry analysis. Biotinylated sAB specifically recognizing bacteria maltose-binding protein (sAB-MBP) was used in a control experiment (**Extended Data Fig. 4a**). Significant hits from the mass spectrometry results with the false discovery rate (FDR) < 0.05 were converted to gene IDs and subjected to gene ontology (GO) analysis. GO enrichment analysis for cellular components showed that K29-linked ubiquitination is present in the spliceosome, proteasome, nuclear body, stress granules, and in particular several important complexes involved in mitosis, including the chromosome, mitotic spindle, and midbody (**Fig. 3c**). GO enrichment analysis for biological processes indicated that the hits were enriched in RNA processing, transcriptional regulation, protein homeostasis, the stress response, and cell cycle regulation (**Fig. 3d-f, Extended Data Fig. 4b**, see supplemental materials for details). The last three categories were further investigated in this work. Specifically, among hits enriched in the protein homeostasis category, VCP (also known as p97 or Cdc48, an ATPase associated with various activity (AAA) involved in the processing of ubiquitinated proteins) and its diverse cofactors, two major autophagic adapters, and many proteasome subunits were identified (**Fig. 3d**). Among hits enriched in the stress response category, stress granule components and some heat shock and oxidative stress responsive proteins were identified (**Fig. 3e**). Among hits enriched in the cell cycle category, proteins that control different stages of the cell cycle, including those involved in DNA replication, spindle assembly, chromatin condensation and midbody formation, were identified (**Fig. 3f**).

### Proteotoxic stress induced K29-linked ubiquitination

To further investigate the results of mass spectrometry and GO analysis, we costained HeLa cells for K29-linked ubiquitin with VCP, the 20S proteasome (the core machinery involved in cellular protein degradation), and eukaryotic translation initiation factor 3 subunit B (EIF3B, enriched in stress granules) using sAB-K29 and the corresponding antibodies. The results showed that K29-linked polyubiquitin was widely distributed in the cytoplasm and nucleus and generally colocalized with VCP and the 20S proteasome (**Fig. 4a, Extended Data Fig. 5a and b**), as well as EIF3B in the cytoplasm (**Fig. 4a**). In addition, K29-linked polyubiquitin formed bright puncta in the cytoplasm of some cells (arrowheads, **Fig. 4a**). It was also found in some large cytoplasmic droplets and costained with VCP and the 20S proteasome, but not EIF3B or K48-linked polyubiquitin (arrows, **Fig. 4a and Extended Data Fig. 5c**).

**Fig. 5.**
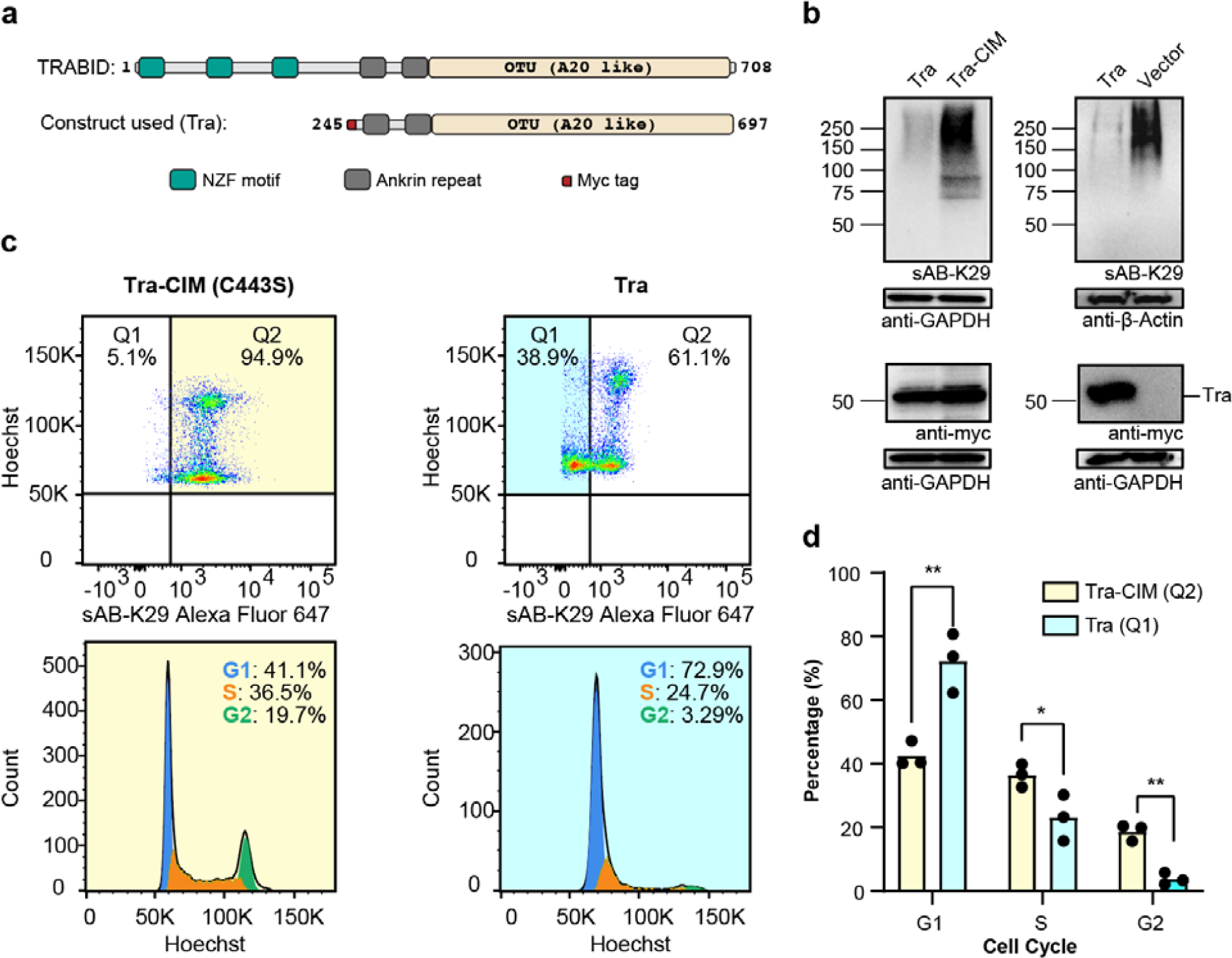
K29-linked polyubiquitination is involved in cell cycle regulation. **a**, A truncated form of the human deubiquitinating enzyme (DUB) TRABID (residues 245-697, named Tra) was used to reduce the level of K29-linked polyubiquitination in HeLa cells. Domain diagrams of full-length TRABID and the Tra construct are shown. **b**, The expression and effect of Tra. HeLa cells were transfected with Tra, Tra-CIM or an empty vector for 24-27 h. The cell lysates were analyzed by Western blotting using a c-myc antibody and sAB-K29, respectively. **c**, Cell cycle analysis of HeLa cells transfected with Tra or Tra-CIM for 24-27 h using flow cytometry. Representative flow cytometry graphs from three biological replicates are shown (a total of 3 injections). Highlighted areas were subjected to cell cycle analysis. **d**, Quantification of the cell cycle analysis results in panel D. Three replicates were included in this experiment. (*, p < 0.05 by two-tailed Student’s *t*-test; **, p < 0.01 by two-tailed Student’s *t*-test). Error bars represent standard deviations.

To further explore this phenomenon, we treated HeLa cells with CuET (bis-(diethyldithiocarbamate)-copper), a drug that targets NPLOC4 ^46, 47^ (also known as Npl4, the most common adapter of VCP); MG132, a proteasome inhibitor; and sodium arsenite, in which CuET and MG132 lead to unfolding protein response and sodium arsenite triggers the stress response. All three compounds dramatically increased the level of K29-linked and K48-linked polyubiquitination in the cell lysate, but only slightly increased K63-linked polyubiquitination, as shown by Western blot analysis (**Fig. 4b and Extended Data Fig. 5d**). The cellular VCP level did not change much upon treatment with CuET or MG132 (**Extended Data Fig. 5j**). We also performed immunofluorescent staining experiments after the treatments. Many more K29-linked polyubiquitin puncta were observed after CuET treatment, and these puncta colocalized with VCP (**Fig. 4c**). Stress granules that stained positive with sAB-K29 were observed after treatment with MG132 or sodium arsenite, which colocalized with the 20S proteasome or EIF3B, respectively (**Fig. 4c**). We further performed these experiments under milder stress conditions, and observed that K29-linked polyubiquitin was still colocalized with VCP and 20S proteasome at a lower concentration of CuET and MG132, respectively (**Extended Data Fig. 5f-g**). However, at a lower concentration of sodium arsenite, much less stress granules were formed and K29-linked polyubiquitin was not colocalized with EIF3B (**Extended Data Fig. 5h**). Finally, we confirmed that the staining was specific, since no signal was observed when a control sAB-MBP was used (**Extended Data Fig. 5e**).

To further investigate the involvement of K29-linked polyubiquitination in the stress response, we treated HeLa cells with heat shock by exposure to 45 °C, which can also lead to stress granule formation, as previously described ^48^. We found that K29-linked polyubiquitin colocalized with both EIF3B and G3BP1 in the resulting stress granules (**Fig. 4d**). G3BP1, another biomarker of cytosolic stress granules ^49^, was identified as a hit from our mass spectrometry results (**Fig. 3e**). Interestingly, we found that stress granules from sodium arsenite treatment were VCP positive, whereas heat shock for 30 minutes resulted in VCP-negative stress granules (**Extended Data Fig. 5i)**. Together, our experiments suggest that K29-linked ubiquitination is upregulated in the proteotoxic stress response and enriched in the granular structures formed after exposure to stress.

### Reduction in the level of K29-linked ubiquitination led to cell cycle arrest

As shown by the results of mass spectrometry and GO analysis, many cell cycle-related proteins and complexes were detected (**Fig. 3c and f**). To further investigate this phenomenon, we designed a method to reduce the level of K29-linked ubiquitination in cultured cells. A reduction in K29-linked ubiquitination was achieved by transfecting the cells with a truncated human DUB, TRABID (residue range 245-697, noted as Tra hereafter). According to previous studies, the Tra construct preferentially targets K29-linked ubiquitin chains, although it shows minimal activity against K33- and K63-linked ubiquitin chains ^43, 50^. As expected, HeLa cells transfected with the Tra construct showed significantly decreased levels of K29-linked polyubiquitinated proteins compared to those cells transfected with catalytic inactive mutant (C443S) of Tra (Tra-CIM) construct and the empty vector, as detected by sAB-K29 (**Fig. 5b and c**). We also tested the level of K63-linked polyubiquitinated proteins and total ubiquitinated proteins using commercially available antibodies. No significant difference between Tra and vector transfection was observed (**Extended Data Fig. 6a-b**).

**Fig. 6.**
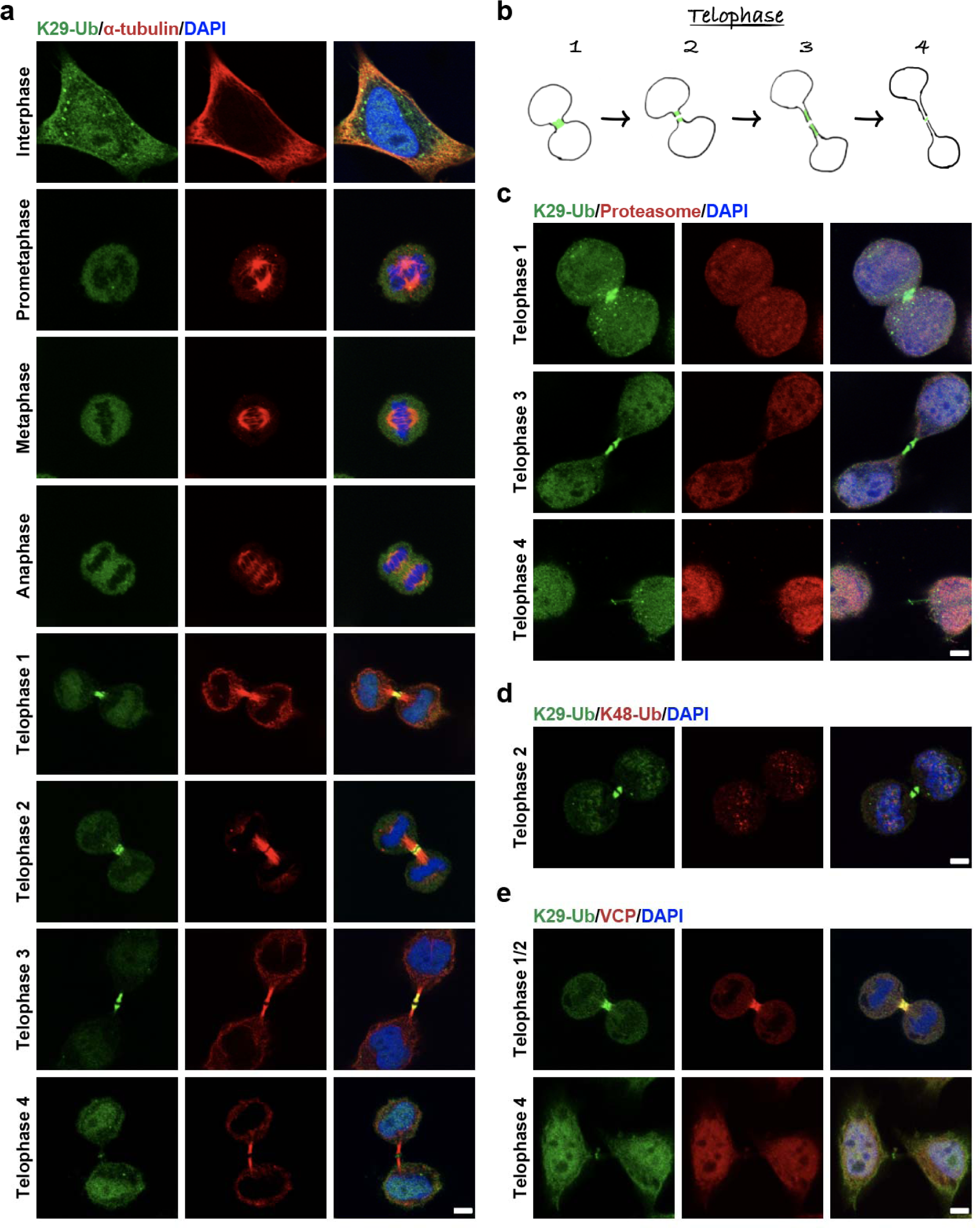
K29-linked polyubiquitination is enriched around the midbody during mitosis. **a**, Immunofluorescent staining of HeLa cells in different stages of the cell cycle. Costaining for K29-Ub and β-tubulin is shown. **b**, A diagram showing migration of the K29-linked polyubiquitination signal during the telophase of mitosis. Roughly four stages could be distinguished based on localization and morphology. **c**, Immunofluorescent staining of HeLa cells at the telophase of mitosis. Costaining for K29-Ub and the proteasome (20S) is shown. **d**, Immunofluorescent staining of HeLa cells at the telophase of mitosis. Costaining for K29-Ub and K48-Ub (Abcam, cat# ab140601) is shown. **e**, Immunofluorescent staining of HeLa cells at the telophase of mitosis. Costaining for K29-Ub and VCP is shown. Scale bars in panels **a**, **c**, **d**, and **e** correspond to 5 μm.

Next, we performed cell cycle analysis of HeLa cells transfected with either the Tra construct or Tra-CIM construct using flow cytometry. Among HeLa cells transfected with the Tra construct, 38.9% of the cells showed a reduced level of K29-linked ubiquitination, as detected by sAB-K29, whereas only 5.1% of the cells transfected with the Tra-CIM were found in the same region (Q1, **Fig. 5d**). The reduced level of K29-linked ubiquitination in the Q1 cells was due to expression of the Tra construct as shown by the signal of cMyc channel (**Fig. 5b and Extended Data Fig. 6c-d**). Cell cycle analysis of Q1 cells transfected with the Tra construct showed that ∼73% of the cells were in G1 phase, ∼24% of cells were in S phase, and only ∼3.2% of cells were in G2 phase, which was significantly different from the cell cycle distribution of Q2 cells transfected with the Tra (**Extended Data Fig. 6f**), Tra-CIM (**Fig. 5c and d**) and normal HeLa cells (**Extended Data Fig. 6e**). In the cell cycle analysis of HeLa cells transfected with either the Tra construct or the empty vector construct, we observed a similar G1/S arrest phenomenon in cells showing reduced level of K29-linked ubiquitination (**Extended Data Fig. 7a and b**). We further tested A549 cells with the same experimental setup and observed similar results (**Extended Data Fig. 7c and d**). Together, these results suggest that K29-linked ubiquitination is essential for cell cycle progression. A reduction in the level of K29-linked ubiquitination arrested the cells in G1 phase.

### K29-linked ubiquitination is involved in midbody assembly

To further investigate the relevance of K29-linked ubiquitination to cell cycle regulation, we performed immunofluorescent staining of HeLa cells at different stages of the cell cycle using sAB-K29 (**Fig. 6a**). As shown by the results of Western blotting, K29-linked ubiquitination occurred in all stages of mitosis, but the level of K29-linked ubiquitination was slightly reduced in cells during mitosis compared to asynchronized cells (**Extended Data Fig. 8a**). Unexpectedly, the K29-linked ubiquitination signal was especially enriched around the midbody in telophase. Based on localization of the K29-linked polyubiquitin signal and the morphology of the daughter cells, we further divided telophase into four stages termed telophase 1-4 (**Fig. 6b**). At telophase 1, the K29-linked polyubiquitin signal formed a bright dot in the middle of two daughter cells. A dark zone appeared at telophase 2. The daughter cells and K29-linked polyubiquitin signal were further apart in telophase 3. Finally, the K29-linked polyubiquitin signal formed a dim ring in the midbody at telophase 4, which was visualized by three-dimensional stochastic optical reconstruction microscopy (3D STORM, **Extended Data Fig. 8b**, **Supplementary Movies 1 and 2**). The positional and morphological changes in the K29-linked polyubiquitin signal during telophase suggest that the modification was dynamic and tightly regulated. We also performed immunofluorescent staining following a pre-extraction protocol to remove weakly bound proteins and found that the K29-linked polyubiquitin signal was still enriched around the midbody at telophase, indicating that K29-linked ubiquitination occurs on strongly interacting or covalently modified midbody components (**Extended Data Fig. 8c and d**).

To test whether K29-linked ubiquitination at the midbody is a degradation signal, we performed immunofluorescent costaining using sAB-K29. K29-linked polyubiquitin colocalized with neither the 20S proteasome (**Fig. 6c**) nor K48-linked polyubiquitin (**Fig. 6d**) around the midbody at telophase, indicating that K29-linked ubiquitination around the midbody may not function as a degradation signal. Previously, VCP was reported to participate in midbody assembly ^51^; therefore, we tested the colocalization of VCP with K29-linked polyubiquitin by costaining and observed their colocalization at telophase 1 and 2 but not late telophase 4 (**Fig. 6e**). Therefore, K29-linked ubiquitination may be a specific signal for VCP-mediated midbody assembly, but this hypothesis requires further investigation.

### Identification of midbody proteins modified with K29-linked polyubiquitin

To identify the substrates of K29-linked ubiquitination in the midbody, we sought to perform immunoprecipitation (IP) under denaturing conditions to minimize the presence of noncovalently bound proteins. We first tested the specificity of sAB-K29 under various denaturing conditions and found that biotinylated sAB-K29 could specifically pull down K29-linked diUb in 1 M urea (**Extended Data Fig. 3e and 9a**). Next, we synchronized HeLa cells to telophase to maximize the abundance of the midbody and lysed the cells using 8 M urea. The lysate was further diluted in 1 M urea, followed by IP using biotinylated sAB-K29. The enriched proteins were separated on an SDS-PAGE gel and subjected to label-free quantitative mass spectrometry analysis.

Several midbody proteins, such as TBK1, PLK1, INCENP, and MKLP1, were enriched with respect to the control (**Extended Data Fig. 9b**). Through immunofluorescent costaining, we observed that TBK1, PLK1 and INCENP colocalized with the K29-linked polyubiquitin signal (top row, **Extended Data Fig. 9c**). Interestingly, MKLP1 did not colocalize with the K29-linked polyubiquitin signal at telophase 2, but with the progression of cytokinesis, it gradually colocalized with the ring structure consisting of K29-linked polyubiquitin formed at telophase 4 (bottom row, **Extended Data Fig. 9c**). We also performed immunofluorescent staining for a well-known midbody regulator, Aurora B, which did not show up as a hit in our mass spectrometry analysis. As expected, the K29-linked polyubiquitin signal colocalized with Aurora B in the midbody (**Extended Data Fig. 9d**), suggesting that sAB-K29 pulled down only K29-linked ubiquitinated substrates under denaturing conditions. We further validated these results by IP and Western blotting. We used antibodies against TBK1, PLK1, INCENP and MLKP1 to pull down proteins in the lysate of HeLa cells synchronized to telophase and used sAB-K29 to detect K29-linked ubiquitination (**Extended Data Fig. 9d**). Alternatively, we used sAB-K29 to pull down proteins in the cell lysate and used antibodies against the respective proteins for Western blotting (**Extended Data Fig. 9e**). As expected, in both experiments, polyubiquitinated substrates with a high molecular weight were observed. In summary, we identified TBK1, PLK1, INCENP and MLKP1 as potential substrates of K29-linked ubiquitination during midbody assembly. The detailed roles of their modification require further investigation.

## DISCUSSION

K29-linked ubiquitination is one of the most abundant polyubiquitin linkage type following K48-linked ubiquitination which serves as a signal for proteasome-mediated degradation. The cellular function of K29-linked ubiquitination is not clear because it is the only linkage type that does not have a specific binder as a tool for detection ^34^. In this study, we filled in this gap by developing and characterizing a linkage-specific sAB against K29-linked polyubiquitin, named sAB-K29, using synthetic K29-linked diUb. We determined the crystal structure of sAB-K29 in complex with recombinant K29-linked diUb and elucidated its binding specificity at the atomic level. Using sAB-K29 as a tool, we discovered that K29-linked ubiquitination is mainly involved in RNA processing, the stress response, and cell cycle regulation. A reduction in K29-linked polyubiquitin by transfection of a DUB construct preferentially hydrolyzing K29 and K33 linkages^43^ resulted in cell cycle arrest. Furthermore, we found that the K29-linked polyubiquitin signal was specifically enriched around the midbody during telophase of mitosis and identified potential substrates of this type of modification.

We explored the role of K29-linked ubiquitination in several stress responses and found that K29-linked ubiquitin was enriched in puncta, liquid droplets, and stress granules in the cytosol upon proteotoxic stress. Our mass spectrometry results identified all the components of proteasome, suggesting a role in protein degradation of this modification. Furthermore, it may also serve as a localization signal since K29-linked ubiquitin did not always colocalize with the 20S proteasome and K48-linked ubiquitin (**Fig. 6c and d**). Moreover, we treated HeLa cells with puromycin, an antibiotic drug that can be incorporated into nascent chains during translation, and found that HeLa cells with a reduced level of K29-linked ubiquitination did not show obvious accumulation of nascent chains (**Extended Data Fig. 10e and f**), which suggested that K29-linked ubiquitination may play other roles besides being a signal for degradation.

We also explored the role of K29-linked ubiquitination in cell cycle regulation. A reduction in K29-linked ubiquitination led to cells arrested in G1/S, but did not affect the cellular levels of some key regulators, including cyclin A, CDC27, STAT3, and cyclin D1 (**Extended Data Fig. 10a-d**). Moreover, the fact that enriched K29-linked ubiquitin did not colocalize with 20S proteasome in the midbody further suggested that K29-linked ubiquitin chains may play nondegradative role in cell cycle regulation. From the mass spectrometry results, several cullin family E3 ligases, including CUL1, CUL3, CUL4A, and CUL4B, may serve as the “writer” of this modification (**Extended Data Fig. 9b**). Further experiments to identify the entire E3 complex responsible for this modification are necessary.

K29-linked ubiquitination was enriched around the midbody during telophase of mitosis. This modification was dynamic, as the localization and structure changed over the progression of mitosis (**Fig. 6a**). We identified multiple midbody proteins that could be substrates of K29-linked ubiquitination from IP experiments under denaturing conditions. The fact that VCP, but not the proteasome, colocalized with the K29-linked polyubiquitin signal again suggested a nondegradative role for this modification. VCP is an ATPase involved in processing various ubiquitinated cellular proteins ^52^. Our results suggested that VCP was widely associated with K29-linked ubiquitination under normal conditions, during stress, or during cell division. Therefore, it may serve as a key processor for this modification. The exact role of K29-linked ubiquitination in midbody assembly requires further investigation.

## Acknowledgements

Funding for this work was, in part, provided by the Catalyst Award from the Chicago Biomedical Consortium. This work was supported by Chicago Biomedical Consortium Catalyst Award C-086 to M.Z. We thank the National Key R&D Program of China (No. 2017YFA0505200), NSFC (91753205) for financial support. We thank Prof. Haiteng Deng, Xianbin Meng and Meng Han in the Proteomics Facility at Technology Center for Protein Sciences, Tsinghua University, for help in mass spectrometry analysis. We thank Vytas Bindokas in the Integrated Light Microscopy Core Facility and David Leclerc in the Flow Cytometry Core Facility at the University of Chicago for help in fluorescent imaging and flow cytometry.

## Author Contributions

Y.Y., M.P., L.L. and M.Z. designed all the experiments and interpreted the results. S.K.E and A.A.K performed the sAB selection and evaluation; Y.Y., Q.Z., and M.P. synthesized all diubiquitin molecules and carried out the related biochemical characterizations. Y.Y., J.L. and M.Z. performed crystal screening and data processing. Q.Z., Y.Y. and M.P. performed and interpreted the LC-MS/MS experiments. Y.Y., Y.X., S.P., and J.F. performed the cell-based imaging experiments. M.Z., M.P., and Y.Y. wrote the paper. M.Z., L.L. and A.A.K supervised the project.

## CONTACT FOR REAGENT AND RESOURCES SHARING

Further information and requests for reagents and resources should be directed to the Lead Contact.

## EXPERIMENTAL MODEL AND SUBJECT DETAILS

### Mammalian cell culture

HeLa cells, 293T cells and A549 cells were cultured in Dulbecco’s Modified Eagle’s Medium (DMEM) supplemented with 10% Fetal Bovine Serum (FBS). Cells were tested for mycoplasma contamination. Plasmids were transfected using lipofectamine 3000 (Invitrogen).

## METHOD DETAILS

### Generation of K29-linked diubiquitin

K29-linked diubiquitin was generated as reported before ^33^. Briefly, 2 mM wildtype ubiquitin was mixed with 1 μM UBA1, 10 μM Ubch7, and 10 μM UBE3C in a buffer containing 50 mM Tris (pH 7.5), 150 mM NaCl, 10 mM ATP, 10 mM MgCl_2_, and 1 mM Tris(2-carboxyethyl)phosphine (TCEP), at 30 °C overnight. The reaction mixture were then incubated with 2 mM vOTU and 5 mM DTT at 30 °C overnight to remove non K29-linked ubiquitin chains. K29-linked diubiquitin was purified by cation exchange using a Source S column (GE Healthcare) equilibrated in 50 mM NaOAc, pH 4.5. The column was eluted with a linear gradient of NaCl up to 1M. The peak corresponding to diubiquitin was pooled and further purified by gel filtration using a Superdex 75 column (GE Healthcare) equilibrated in 25 mM Tris, pH 8.0, 150 mM NaCl and 0.5 mM TCEP.

### Generation of polyubiquitin chains

Polyubiquitin chains were generated as reported before ^53^. Briefly, wildtype ubiquitin or K29C mutant was mixed with UBA1 and specific combinations of E2 and E3 enzymes in a buffer containing 20 mMTris, pH 7.5, 150 mM NaCl, 10 mM ATP, 10 mM MgCl_2_, and 1 mM TCEP. Specifically Ubch7/UBE3C were used to generate a mixture of polyubiquitin containing K29-linked chains; Ube2s, Ube25K, and Ubc13/Mms2 were used to generate K11-, K48- and K63-linked polyubiquitin chains, respectively. The reactions were monitored by Western Blot.

### Synthesis of biotinylated K29-linked diUb

Biotinylated K29-linked diUb was synthesized using hydrazide-based native chemical ligation and microwave-assisted solid phase peptide synthesis (SPPS) as reported before^54^. Briefly, biotinylated K29-linked diUb was divided into three peptide segments, namely fragment **1** Ub[Met_1_-Phe_45_]-NHNH_2_, fragment **2** Ub[Cys_46_-Gly_76_]-NH2, and fragment **3** Ub [Cys(Acm)_46_-Gly_75_]-G-[Met_1_-Lys_29_-Phe_45_]-NHNH_2_ (**Extended Data Fig. 1**). Ala46 of each Ub subunit was temporarily mutated to Cys for native chemical ligation. The isopeptide bond at Lys29 of fragment **3** was constructed using the orthogonal building block Fmoc-Lys(Alloc)-OH rather than normal Fomc-Lys(Boc)-OH^55^. An acetamidomethyl (Acm) group was used to protect the thiol group of N-terminal Cys of to avoid oligomerization or self-cyclization. All the peptides were synthesized using standard Fmoc SPPS protocols under microwave conditions (CEM Liberty Blue). The insertion of PEG linker was constructed using the building block Fmoc-mini-PEG, and biotin was directly condensed to the N termini. Fragment **4** was synthesized by hydrazide-based native chemical ligation of fragment **1** with **2** followed by Acm deprotection. Finally, fragment **2** and fragment **4** were ligated to generate the full-length diUb, which was subjected to desulfurization to convert Cys46 in the two Ub units into Ala46.

### Purification of synthetic antibody fragment (sAB)

Plasmids containing sAB-K29 or sAB-MBP were transformed to E. coli BL21(DE3) cells for over-expression. Cells were collected and lysed in the Lysis Buffer (20 mM Tris-HCl, pH 7.4, 500 mM NaCl, 1 mM phenylmethylsulfonyl fluoride (PMSF)). The cell lysate was cleared by centrifugation and run through a protein G-sepharose column equilibrated in the lysis buffer. The sABs were eluted using the Elution Buffer (0.1 M glycine, pH 2.7) and then cation-exchanged using a Mono S column equilibrated in Buffer A (50 mM sodium acetate pH 5.0). The proteins were eluted using a linear gradient of the sodium ions by mixing with Buffer B (50 mM sodium acetate pH 5.0, 2M NaCl).

### Purification of other proteins

Plasmids containing UBA1 (homo sapiens), UBCH7 (homo sapiens), vOTU (the viral ovarian tumor DUB encoded by Crimean Congo hemorrhagic fever virus) were transformed to E. coli BL21(DE3) cells. UBA1 was over-expressed by IPTG induction, UBCH7 and vOTU were over-expressed by autoinduction following the established protocol ^56^. UBE3C was transformed to E. coli Rosseta 2 (DE3) cells and over-expressed by IPTG induction. The purification procedures followed the established protocols ^53, 57^. Briefly, *E. coli* cell pellets were resuspended in Lysis Buffer containing 50 mM Tris, pH 8.0, 300 mM KCl, 20 mM imidazole and 1 mM DTT with 1mM PMSF added freshly. Subsequently, cells were lysed by sonication, and the lysates were cleared by centrifugation at 16500 rpm for 30 min. Then the supernatants were flowed through a Ni–NTA gravity column twice at 4 °C. Beads were washed with Wash Buffer containing 50 mM Tris, pH 8, 150 mM NaCl and 25 mM imidazole. Finally, the His-tagged proteins were eluted in Elution Buffer containing 50 mM Tris, pH 8.0, 300 mM NaCl, 400 mM imidazole and 1mM DTT. For His-SUMO-Ube3c, SUMO protease (Ulp1p) were added to the eluate, and dialyzed in the buffer containing 25 mM Tris, pH 8.0, 150 mM NaCl and 0.5 mM TCEP. The proteins were further purified by a Superdex 200 gel filtration column (GE Healthcare) equilibrated in the buffer containing 25 mM Tris, pH 8.0, 150 mM NaCl and 0.5 mM TCEP.

### Crystallization of sAB-K29 and K29-linked diubiquitin complex

Complex of sAB-K29 and K29-linked diubiquitin was formed by mixing purified sAB-K29 and K29-linked diubiquitin (1:1.5), followed by size exclusion chromatography using a Superdex 75 column equilibrated in a buffer containing 20 mM Tris-HCl, pH 7.4, 150 mM NaCl. Crystals were grown using the hanging drop method and appeared in 1 weeks at 21 °C from a 1:1 mixture of protein (18 mg/mL) and reservoir solution (0.1 M phosphate-citriate, 40% v/v Polyethylene glycol 300). Crystals were harvested and flash-frozen in liquid nitrogen without any cryo-protectant.

### Data collection and structure determination

X-ray diffraction data were collected at the Northeastern Collaborative Access Team beamline (24-ID-E) at the Advanced Photon Source (Supplemental Table 1). The diffraction data were integrated and scaled using XDS/XSCALE ^58^. The structure was determined by molecular replacement using PHASER ^59^ in PHENIX ^60^ and a Fab search model. The model was built using COOT ^61^ and was refined using PHENIX. Coordinates and structure factor amplitudes have been deposited in the Protein Data Bank under accession number 7KEO.

### Western blot

Western blots were performed using the standard protocol. Peroxidase-conjugated goat anti-human IgG, F(ab)2 fragment specific (Jackson ImmunoResearch, 109-036-006) was used as the secondary antibody (1:5000) when using sAB-K29 as the primary binder (0.2-2 μg/mL). Other primary antibodies are diluted as followed: Anti K63-Ub (Sigma,1:1000); Anti Ub (Abcam, 1:1000); Anti β-Actin (Santa Cruz, 1:1000), Anti GAPDH (Santa Cruz, 1:2000), Anti c-Myc (Genscript, 0.3 μg/mL), Anti VCP (Santa Cruz, 1:1000), Anti INCENP (Abcam,1:3000), Anti MKLP1 (Abcam,1:1000), Anti TBK1 (CST,1:1000), Anti PLK1 (Abcam, 1 μg/mL).

### Phage Display

Four rounds of biopanning using Fab Library E (kind gift of A. Koide) were performed as follows using a protocol adapted from previous studies ^41, 42^. The first round of selection was performed manually using 300 nM of biotinylated K29-linked diubiquitin immobilized onto 250 µL of Streptavidin Paramagnetic Particles (Promega). Following washing with Selection Buffer (SB; 20 mM Hepes, pH 7.5, 150 mM NaCl, 0.5% BSA, 0.05% Tween-20), the beads were incubated with Library E for one hour at room temperature (RT). The beads were then washed three times with SB and used to directly infect log-phase *E. coli* XL-1 blue cells. Following infection, 2XYT media supplemented with ampicillin and M13 helper phage was added and phage were amplified overnight. To increase the stringency of selection, three additional rounds of sorting were performed in the presence of 1.5-µM of mono-ubiquitin as a competitor in solution. Further, the concentration of K29-linked diubiquitin was reduced each round as follows: 150 nM in round 2, 75 nM in round 3, and 15 nM in round 4. These three rounds were performed at RT semi-automatically using a Kingfisher magnetic beads handler. The overnight-amplified elution pool from each preceding round of selection was used as the input for the following round of sorting.

### Single-point competitive phage ELISA

Elution pools from rounds 3 and 4 were used to infect log-phase *E. coli* XL-1 blue cells and plated onto LB-agar supplemented with ampicillin. Twenty-four colonies from each pool were picked and sequenced at the University of Chicago Comprehensive Cancer Center DNA Sequencing facility. DNA sequencing determined that 47 out of 48 colonies corresponded to a single clone, and both clones were tested for binding to K29-linked diubiquitin by ELISA. All ELISA were performed in 96-well plates (Nunc) coated with 2 µg/mL Neutravidin and blocked with phosphate-buffered saline containing 1% BSA. The plates were washed with ELISA Buffer (EB; 25 mM Hepes, pH 7.4, 150 mM NaCl, 0.05% Tween-20) and 25 nM of K29-linked Di-Ub (in EB) was immobilized onto the plates. Amplified phage for both clones were diluted 20X into EB either in the absence or presence of competitors: non-biotinylated K29-linked diubiquitin, mono-ubiquitin, K27-linked diubiquitin, and K33-linked diubiquitin. Phage dilutions with and without competitors were added to target-immobilized ELISA plates, allowed to bind for 15 minutes, washed, and then incubated with an anti-M13 HRP-conjugated secondary antibody (GE Healthcare) for 30 minutes. Following washing with EB, bound phage were detected using TMB substrate kit (Thermo). Specificity was determined by comparing the ELISA signal with and without competition. Each assay was performed in triplicate. Based on ELISA signal, only the most abundant clone from sequencing was found to be specific to K29-linked diubiquitin. This Fab, named sAB-K29, was then subcloned into the expression vector RH2.2 (kind gift of S. Sidhu) using the In-Fusion Cloning Kit (Takara) and sequence-verified.

### Characterization of binding affinity

The binding of sAB-K29 to K29-linked diubiquitin was verified by multi-point ELISA ^62, 63^ and surface plasmon resonance (SPR). ELISA were performed using plates prepared as above. Biotinylated constructs of K29-, K27-, and K33-linked diubiquitin were immobilized individually onto ELISA plates at 25 nM final concentrations. Twelve serial dilutions of sAB-K29, starting at 1 µM concentration, were prepared in EB and each Fab dilution was bound to wells containing either immobilized targets or no targets (to determine any background signal). Following washing with EB, bound Fab was detected by incubation with an anti-Fab HRP-conjugated secondary (Jackson) and TMB substrate kit, as above. The quenched absorbance signal was measured and plotted as a function of the log of Fab concentration. EC_50_ values were then calculated in GraphPad using a variable slope model and assuming sigmoidal dose response. Each assay was performed in triplicate and the mean signal was used to fit the dose response curve.

SPR experiments were performed using a MASS-1 instrument (Bruker) with a His-capture sensor chip (XanTec). His-tagged constructs of K29, K27, and K33-linked diubiquitin were immobilized onto sensor chips following coating of the experimental and reference channels with 0.5 mM NiSO_4_. His-tagged proteins (ligands) were immobilized only in the experimental channels. After optimization of the amount of immobilized ligands based on response units (RUs), a series of sAB-K29 concentrations was injected into both the experimental and reference channels. sAB-K29 was injected over a 60s-association phase, followed by a 600s dissociation time ^63^. Following each injection, the chip surface was regenerated using 350 mM EDTA and 50 mM NaOH, and ligands were subsequently immobilized for the following injection. For each ligand, all experiments were performed on single channels for consistency. Fab injections were then double-reference subtracted using the reference channel signal and a buffer-only experimental channel injection. The double-reference subtracted curves were then fit with one-to-one Langmuir binding models in Scrubber to determine kinetic binding parameters.

### Generation of biotinylated sAB-K29 using sortase enzyme

LPETGG-His_6_ tag was introduced to the C terminus of the heavy chain of sAB-K29. Purified sAB-K29-LPETGG-His_6_ (20 µM) and synthesized biotin peptide (biotin-AEEA-GGGGG, 100µM) were incubated with His-tagged sortase (10 µM) in sortase buffer (10 mM CaCl_2_, 50 mM HEPES, 150 mM NaCl, pH 7.4) for 30 minutes at room temperature. The reaction mixture was then incubated with Ni-NTA beads to remove the remaining His-tagged sAB-K29 and sortase. Biotinylated sAB-K29 in flow through was further purified using Protein A column to remove the biotin peptide.

### Immunoprecipitation for mass spectrometry

Biotinylated sAB-K29 (100 µg) was incubated with 1 mL streptavidin-magnetic beads (Promega, Z5482) prewashed three times with PBS buffer in a thermo shaker for 3 h at 25 °C. The beads containing immobilized sAB-K29 were then incubated with 5 mg of HeLa cell lysate in RIPA lysis buffer (P0013B, Beyotime) on a rotation wheel at 4 °C overnight. The beads were washed 3 times with the wash buffer containing 25 mM Tris-HCl, pH 8.0, 500 mM NaCl, and 0.5 % Triton, followed by 3 times with the wash buffer without Triton. The bound proteins were eluted with 100 µl 8M Urea for 1h at 25 °C in a thermo shaker. Similar procedures were followed for immunoprecipitation under the denaturing condition (denature IP), except that 5 mg of synchronized HeLa cell lysate in 8 M Urea with protease inhibitor cocktail (Roche, 4693116001) was diluted by PBS buffer to a final urea concentration of 1 M before incubating with the conjugated beads. Each experiment was performed in triplicate and eluted fractions were pooled and analyzed by SDS-PAGE or Western blot.

### Sample preparation for label-free mass spectrometry

Eluted protein fractions by denature IP were incubated in 20 µl 4 × LDS loading buffer (NP0007, Thermo Fisher) for 5 min at 95 °C. Proteins were separated on 4%-12% gradient gels (NP0336Box, Thermo Fisher) and stained with Coomassie blue R-250 (161-0436, Bio-Rad). For in-gel digestion, each gel lane was cut into 6 bands which were then cut into smaller (around 1 cm) fragments. Gel pieces were de-stained in 50 mM NH_4_HCO_3_/ACN (1:1, v/v). After washing with 50 mM NH_4_HCO_3_, proteins were reduced by the addition of 25 mM DTT in 50 mM NH_4_HCO_3_ for 60 min at 55 °C followed by alkylation in 55 mM iodoacetamide in 50 mM NH_4_HCO_3_ for 60 min at 37 °C. After washing in 50 mM NH_4_HCO_3_/ACN (1:1, v/v) and dehydration in ACN, proteins were digested overnight at 37 °C with trypsin (1:50, w/w) (V5280, Promega). Peptides were extracted from the gel in 50% ACN/0.1% formic acid, followed by vacuum drying and re-dissolved by 20 µl 0.1 % formic acid for LC-MS analysis.

### Mass spectrometry data collection

Tryptic peptides were separated on an UltiMate 3000 RSLCnano System (Thermo Scientific, USA) at a flow rate of 300 nL/min using a 120 min gradient from 5% ACN, 0.1% formic acid to 45% ACN, 0.1% formic acid followed by a washing step at 100% ACN. Each of the 3 independent biological replicates was measured as technical duplicates. Mass spectra were collected on a Thermo Orbitrap Fusion Lumos mass spectrometer (Thermo Fisher Scientific). Each analysis was operated in data dependent top-speed mode with dynamic exclusion set at 30 s and a total cycle time of 3 s. Full scan MS spectra were acquired in the Orbitrap at a resolution of 120000 (at m/z 200) with an automatic gain control ion target value of 4e5 and a maximum injection time of 50 ms. The most intense precursors with charge states of 2-7 and intensities greater than 5e3 were selected for MS/MS experiments using higher-energy collisional dissociation (HCD) with 27 % collision energy. Isolation was performed in the quadrupole with an automatic gain control ion target value of 2e3 and a maximum injection time of 300 ms.

### Mass spectrometry data analysis and quantification

Raw data from LC-MS/MS measurements were analyzed using MaxQuant version 1.6.5.0 with default settings and label-free quantification (LFQ) enabled. For protein identification, the human reference proteome downloaded from the UniProt database (download date: 2019-07-01) and an integrated database of common contaminants was used. Identified proteins were filtered for reverse hits and common contaminants at a false discovery rate of 5%. Further data processing was performed using Perseus software (version 1.5.8.5) and R software (version 3.6.1). The mass spectrometry proteomics data have been deposited to the ProteomeXchange Consortium via the PRIDE partner repository with the dataset identifier PXD024428 and PSD0244425^64^.

### Immunofluorescence staining and imaging

HeLa cells were fixed with 4% paraformaldehyde in PBS at room temperature for 20 min, washed three times with PBS. Nonspecific binding sites were blocked with 10% normal goat serum, 0.1% Triton X-100 for 1 h at room temperature. The cells were then stained for 2 h at room temperature with the primary binder / antibody. After that, the cells were washed three times with PBS containing 0.1% TritonX-100, following by staining for 1 h with the secondary antibody. Finally, cells were washed three times with PBS containing 0.1% Triton X-100, and mounted with ProLong Gold Antifade reagent with DAPI (Invitrogen). Alexa Fluor 488 or 647 conjugated F(ab’) fragment goat anti-human IgG, F(ab’)2 fragment specific (Jackson ImmunoResearch) was used as the secondary antibody (1:500) when using sAB-K29 as the primary binder (2 μg/mL). Other primary antibodies are diluted as followed: Anti AuroraB (Fisher,1:500); Anti α-Tubulin (Genscript, 1:50-1:100); Anti α-Tubulin (proteintech, 1:200); Anti c-Myc (Genscript, 0.5 μg/mL); Anti EIF3B (Santa Cruz, 1:100); Anti INCENP (Santa Cruz,1:100), Anti MKLP1(Santa Cruz, 1:100); Anti Proteasome 20S core subunits (Fisher, 1:250); Anti K48-Ub (Abcam, 1:200); Anti VCP (Santa Cruz,1:200); Anti G3BP1 (proteintech,1:500); Anti pTBK1 (cell signaling,1:50); Anti PLK1 (Santa Cruz,1:50). Images were collected by a Leica SP5 2-photon laser scanning confocal microscope.

### Super-resolution microscopy

sAB-K29 was labeled with Alexa Fluor 647 (A647, Invitrogen A20006) following the instructions of Alexa Fluor Protein Labeling Kits (Thermo Fisher). Unreacted free dye was removed using P-6 Gel Columns (Bio-Rad). The labeling efficiency is about 1 Alexa Fluor 647 dye per antibody. 3D STORM imaging was conducted as previously described ^65^ on an inverted microscope using a 647 nm laser and a 405 nm laser and data were recorded by an EMCCD camera. For 3D imaging, a cylindrical lens was placed in the emission beam path. Each STORM image was reconstructed from about 5,000 image frames. Imaging buffer was composed of 10 mM NaCl, 50 mM Tris (pH 8), 10% glucose, 50 U/mL glucose oxidase (Sigma Aldrich G2133-10KU), 404 U/mL catalase (EMD Millipore 219001) and 20mM cysteamine (Sigma Aldrich 30070-10G). 3D Image reconstruction and visualization was conducted as previously described ^66^.

### Cellular stress assays

HeLa cells were treated with CuET (at 1 μΜ or 0.5 μM, 4h, this paper), MG132 (20 μΜ or 10 μM, 4h, Sigma-Aldrich) NaAsO_2_ (500 μM, 1h or 250 μM, 4h, Sigma), Cycloheximide (100 μg/ml, 6h, Sigma), Puromycine (1 μg/ml,2h, GoldBio), α-Amanitin (5 μg/ml, 24h, MCE), or 45 °C heat shock for 30 min. K29-ubquitin chains was detected by western blot or immunofluorescence microscopy.

### Cell synchronization

HeLa cells were first synchronized in G1/S-phase with 2 mM thymidine for 24h, followed by PBS wash and releasing to fresh DMEM/10% FBS media for 3h. After that, cells were treated with 50 ng/ml nocodazole for 12h and arrested in prometaphase. For progression through mitosis, cells were harvested via pipetting, washed with PBS twice, and released into fresh media. Cell lysates were taken at indicated time points and analyzed via western blot.

### Flow cytometry

Cells were transfected with the Tra construct (human TRABID, 245-697), the Tra-CIM (human TRABID, 245-697, C443S) or pCMV vector using lipofectamine 3000 for 24-27 h before harvesting and washing with PBS. The cells were fixed with either 1% or 4% PFA for 15 min at room temperature, followed by washing with PBS and permeabilization with 70% ethanol at 4 °C overnight. Nonspecific binding was blocked with 2% BSA. The cells were then incubated with Anti-c-myc antibody conjugated with PE (sc-40, Santa Cruz Biotechnology, 1:40) and either sAB-K29 as primary antibody (3 ug/mL), and Alexa Fluor 647 conjugated F(ab’) fragment goat anti-human IgG, F(ab’)2 fragment specific (Jackson ImmunoResearch) as secondary antibody, or sAB-K29 directly labeled with Alexa Fluor 647 for 1h at room temperature, followed by washing with PBS. Finally, the cells were incubated with Hoechst dye at room temperature for 15 min, followed by washing with PBS. The data was acquired by a LSRFortessa X-20 cell analyzer (BD) and analysis by flowjo (10.6.2).

## SUPPLEMENTAL TABLES

**Extended Data Table 1.**
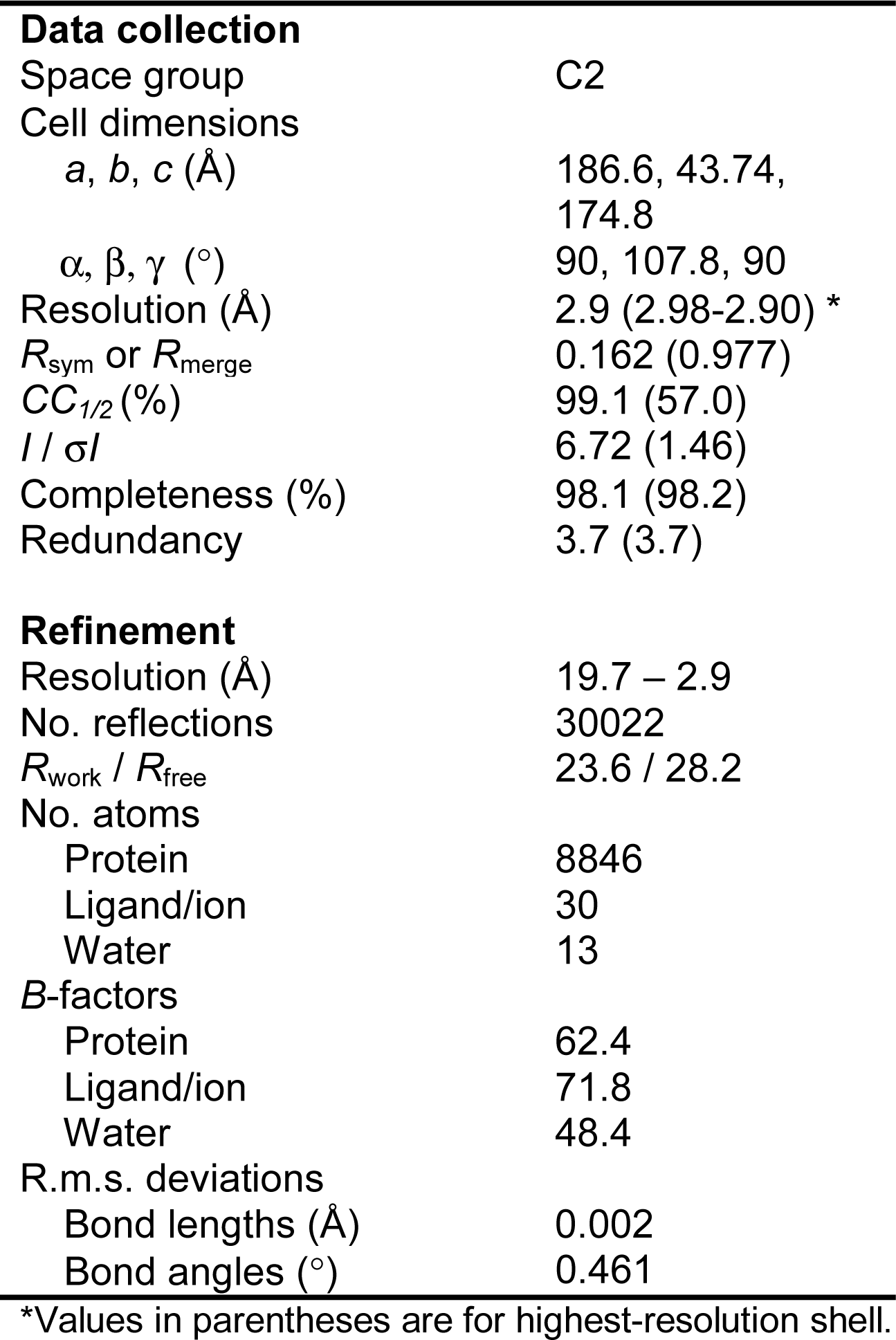
X-ray data collection and refinement statistics.

## SUPPLEMENTAL FIGURE LEGENDS

**Extended Data Fig. 1.**
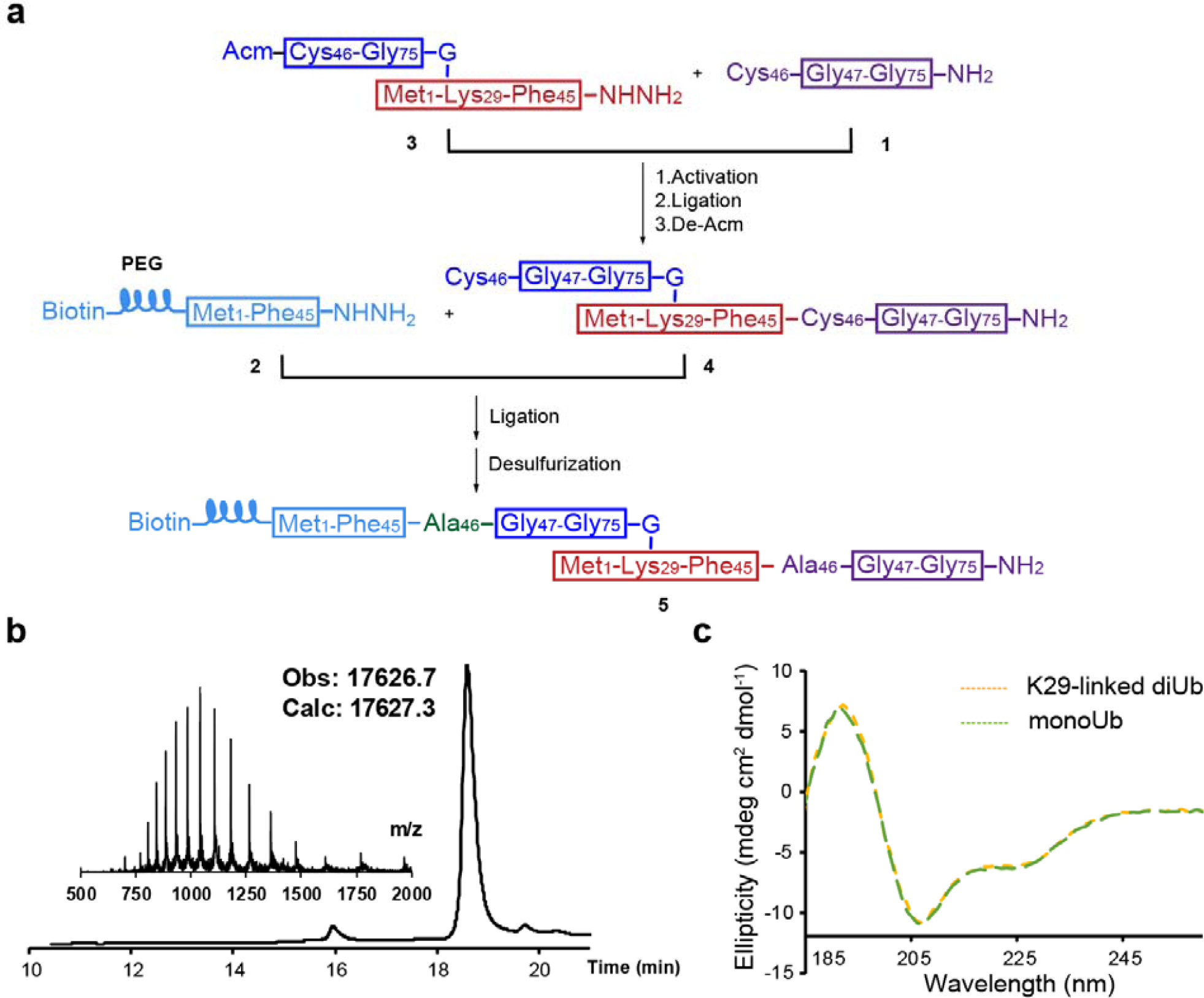
Chemical synthesis and characterization of biotinylated K29-linked diubiquitin. **a**, Synthetic routes for biotinylated K29-linked diubiquitin. **b**, Liquid chromatography and mass spectrometry (LC-MS) analysis of synthetic biotinylated K29-linked diubiquitin. **c**, Circular dichroism (CD) spectra of synthetic biotinylated K29-linked diubiquitin and recombinant monoubiquitin.

**Extended Data Fig. 2.**
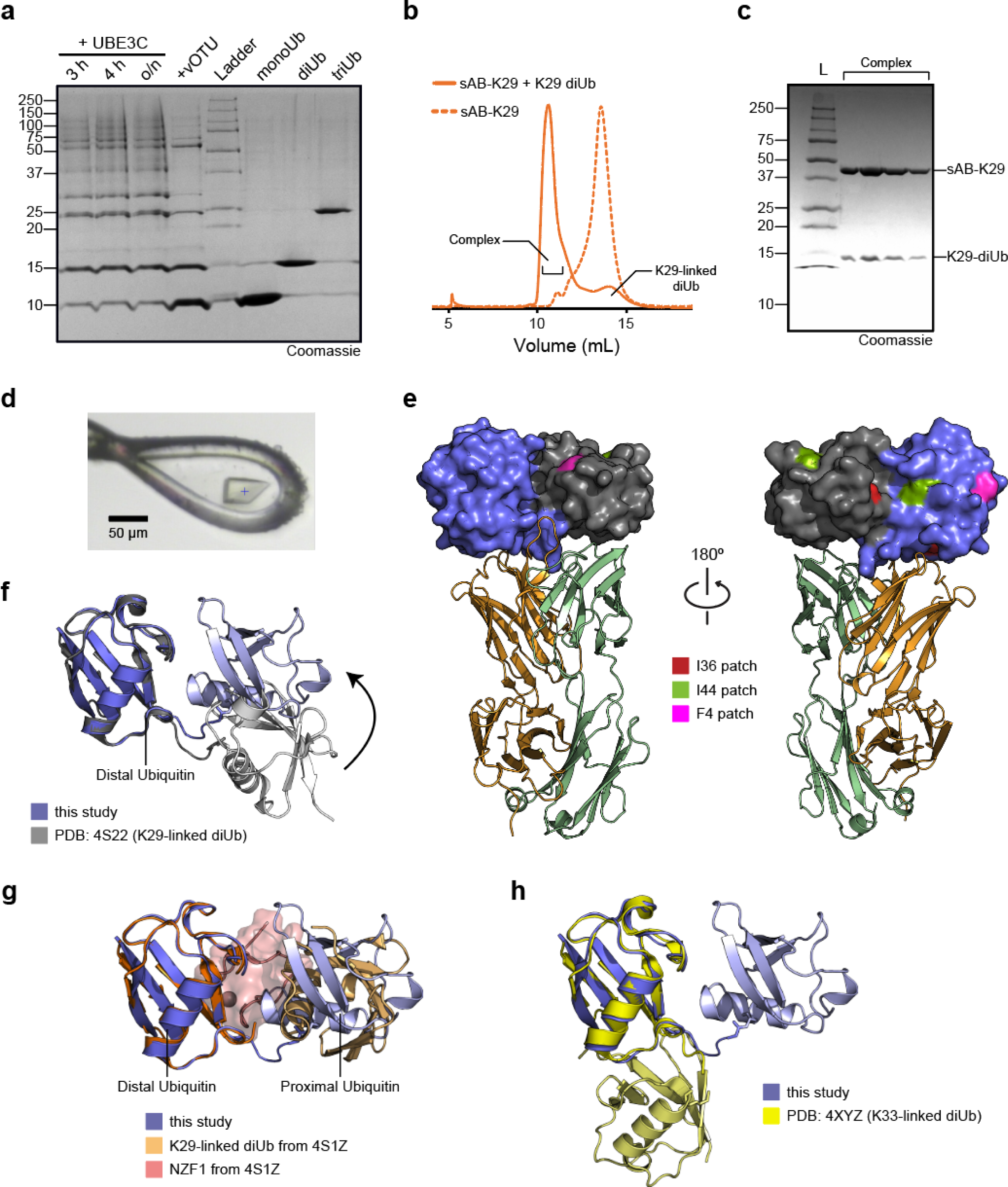
Purification and crystallization of the complex between K29-linked diubiquitin and sAB-K29. **a**, Purification of K29-linked diubiquitin. A mixture of K29- and K48-linked polyubiquitin mixture was assembled by incubating monoubiquitin, UBE1, UBE2L3, and UBE3C overnight. vOTU, a DUB that does not cleave K29-linked chains, was added to the reaction mixture to cleave K48-linked polyubiquitin chains. The resulting mono-, di-, and triubiquitin molecules were separated by anion exchange chromatography. **b**, Size exclusion chromatography (SEC) of sAB-K29 and sAB-K29 in complex with K29-linked diubiquitin. **c**, The SDS-PAGE gel of the complex peak in panel B. **d**, A picture of the crystal of the complex mounted in a cryo-loop. **e**, The distribution of hydrophobic patches on K29-linked diUb. Only the I36 patch of the distal ubiquitin molecule is involved in the interaction with the heavy chain of sAB-K29. **f**, Superimposition of the crystal structure of K29-linked diUb (PDB accession code: 4S22) and K29-linked diUb in the complex structure determined in this study. The distal ubiquitin molecules in the two structures were aligned. **g**, Superimposition of the crystal structure of K29-linked diUb in complex with NZF1 domain of TRABID (PDB accession code: 4S1Z) and K29-linked diUb in the complex structure determined in this study. The distal ubiquitin molecules in the two structures were aligned. NZF1 domain was colored in pink. **h**, Superimposition of the crystal structure of K33-linked diUb (PDB accession code: 4XYZ) and K29-linked diUb in the complex structure determined in this study. The distal ubiquitin molecules in the two structures were aligned. The K29-linked diUb from this study in panels **f-h** are in the same orientation as in Fig. 1c and left of panel **e**.

**Extended Data Fig. 3.**
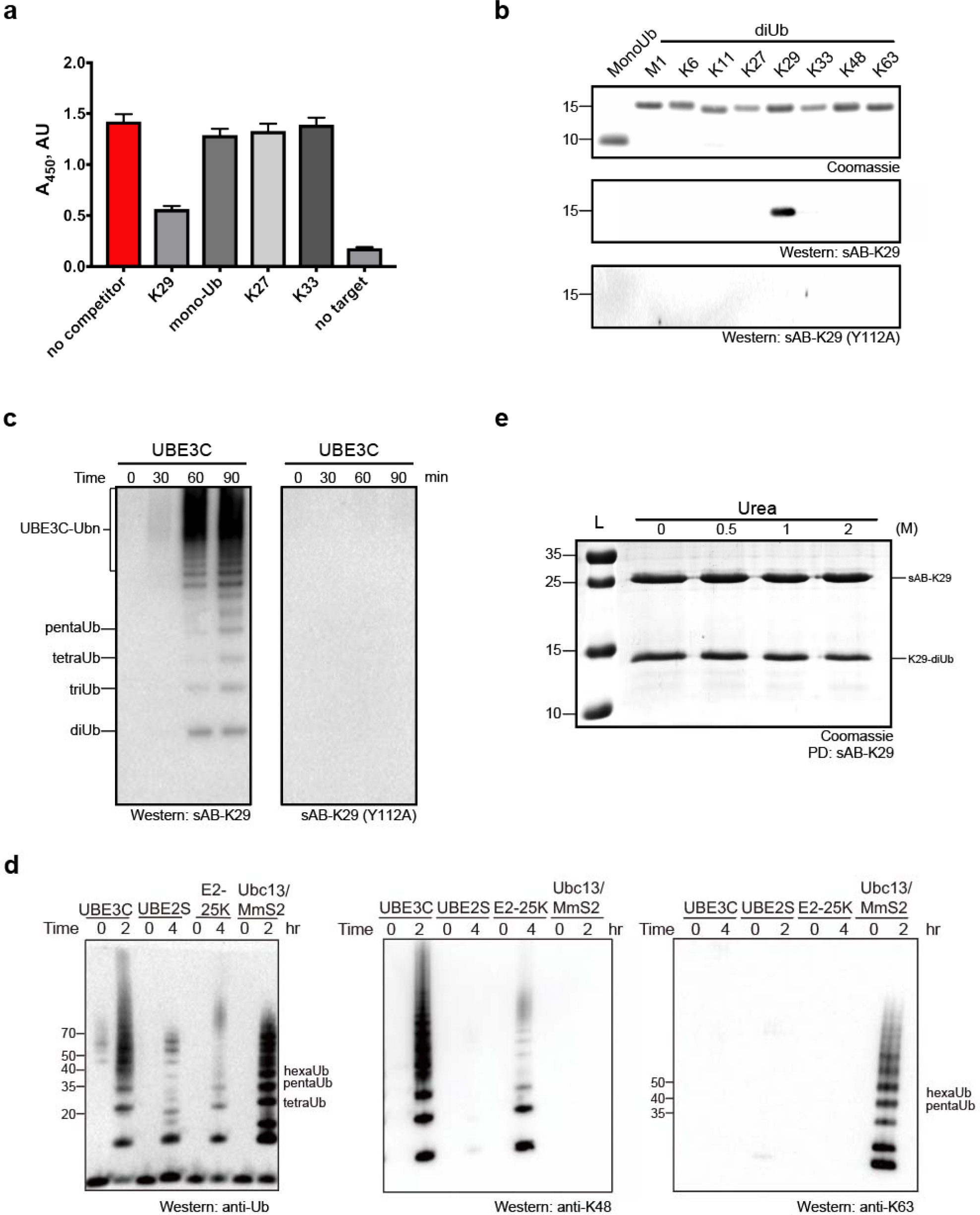
Characterization of sAB-K29 specificity. **a**, Single-point competitive phage ELISA. Biotinylated K29-linked diubiquitin was immobilized and incubated with phage-displayed sAB-K29 in the presence of excess competitors (monoubiquitin and K29-, K27-, and K33-linked diubiquitin) in solution. A decrease in absorption indicated more specific binding. Error bars represent the standard deviations of triplicate experiments. **b**, A point mutation in the heavy chain of sAB-K29 (Y112A) abolished its recognition of K29-linked diUb. Monoubiquitin and diubiquitin (∼500 ng) with eight types of linkages were loaded on an SDS-PAGE gel. Western blotting was performed using sAB-K29 or sAB-K29 (Y112A) as the primary binder. **c**, Mutation of the heavy chain of sAB-K29 (Y112A) abolished its ability to detect K29-linked polyubiquitin chains by Western blotting. K29-linked polyubiquitin chains were assembled by an E1-E2-E3 mixture containing UBE1, UBE2L3, and UBE3C. The reactions were followed for 90 minutes at four time points. **d**, Polyubiquitin chains were assembled by four ubiquitination systems: UBE3C (K48- and K29-linked chains), UBE2S (K11-linked chains), E2-25K (K48-linked chains), and UBC13/Mms2 (K63-linked chains). Parallel Western blotting using anti-ubiquitin antibody, anti-K48 linked ubiquitin chain and K63-linked ubiquitin chain were performed for comparison. **e**, sAB-K29 could be used to pull down K29-linked diUb in vitro under denaturing conditions (up to 2 M urea).

**Extended Data Fig. 4.**
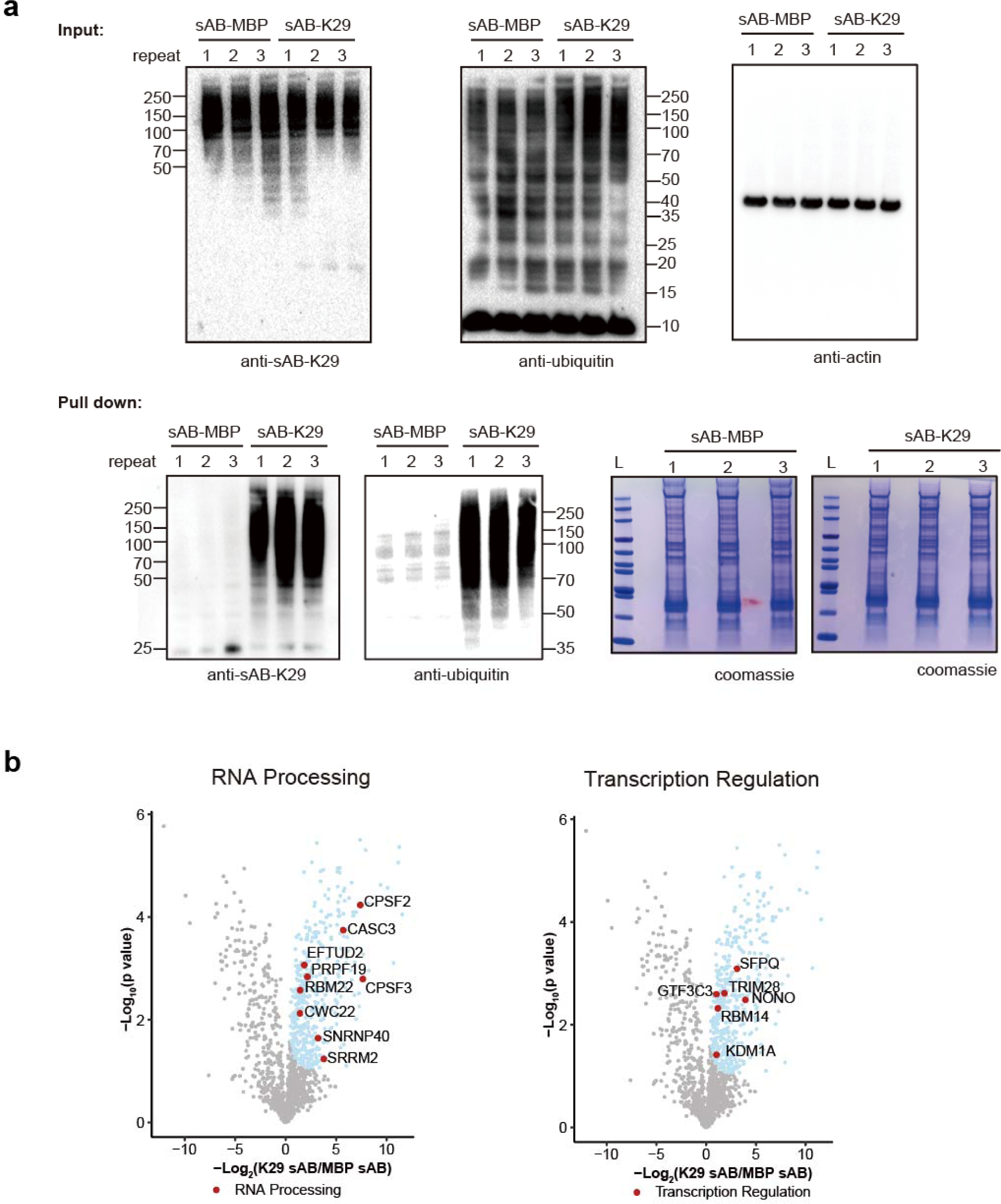
Pull-down and proteomic analysis of HeLa cells using sAB-K29. **a**, Pull-down assay of HeLa cell lysate using sAB-K29. The bound proteins were separated on an SDS-PAGE gel. Gel slices were subjected to label-free quantitative mass spectrometry to identify the enriched proteins. Three biological replicates were performed. sAB-MBP was used in the control pull-down assay. Western blot analysis of the pull-down eluant using sAB-K29 or anti-ubiquitin antibody as the primary binder confirmed that the sAB-MBP is an effective control. **b**, Volcano plots of the quantitative mass spectrometry results. Significantly enriched hits (FDR < 0.05) are colored cyan, with some well-documented proteins involved in RNA processing and transcription regulation highlighted and labeled.

**Extended Data Fig. 5.**
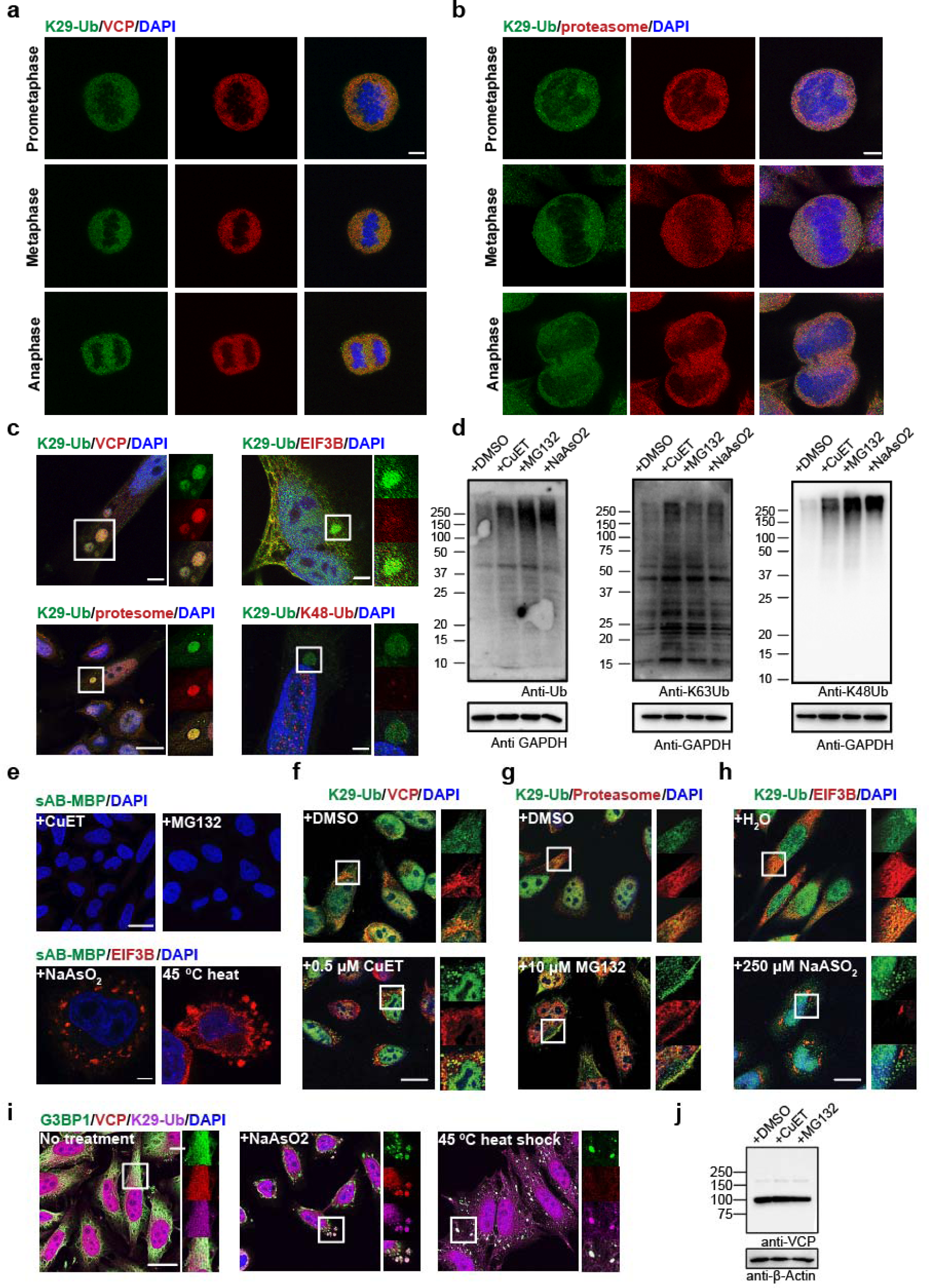
K29-linked ubiquitination is involved in protein homeostasis and the stress response. **a-b**, Immunofluorescent staining of HeLa cells at prometaphase, metaphase, and anaphase of mitosis. Costaining for K29-Ub with VCP (panel A) or the proteasome (20S, panel B) is shown. Scale bars correspond to 5 µm. **c**, K29-linked ubiquitination is enriched in liquid droplets occasionally observed in normal HeLa cells. Costaining for K29-Ub with VCP, EIF3B, the proteasome (20S), and K48-Ub is shown. **d**, Western blots of total ubiquitin, K63- and K48-linked ubiquitin in HeLa cells treated with CuET (an inhibitor of the VCP cofactor Npl4, 1 μM for 4 h), MG132 (a proteasome inhibitor, 20 μM MG132 for 4 h), or sodium arsenite (an oxidative stress inducer, 500 μM for 1 h). **e**, Immunofluorescent staining of HeLa cells with a control sAB (against maltose-binding protein) and costaining for EIF3B in HeLa cells treated with 500 µM sodium arsenite for 1 h or subjected to heat shock at 45 °C for 30 minutes. Scale bars in the top and bottom panels correspond to 20 and 5 μm, respectively. **f-h**, Immunofluorescent staining of HeLa cells treated with 0.5 μM CuET for 4 h, 10 μM MG132 for 4 h, or 250 μM sodium arsenite for 4 h. Costaining for K29-Ub with VCP, the proteasome (20S), and EIF3B is shown. **i**, Immunofluorescent staining of HeLa cells treated with 500 μM sodium arsenite for 1 h or subjected to heat shock at 45 °C for 30 minutes. Costaining for K29-Ub with G3BP1 and VCP is shown. The scale bar corresponds to 20 μm. **j**, Western blot of VCP in HeLa cells treated with CuET (1 μM for 4 h) and MG132 (20 μM for 4 h).

**Extended Data Fig. 6.**
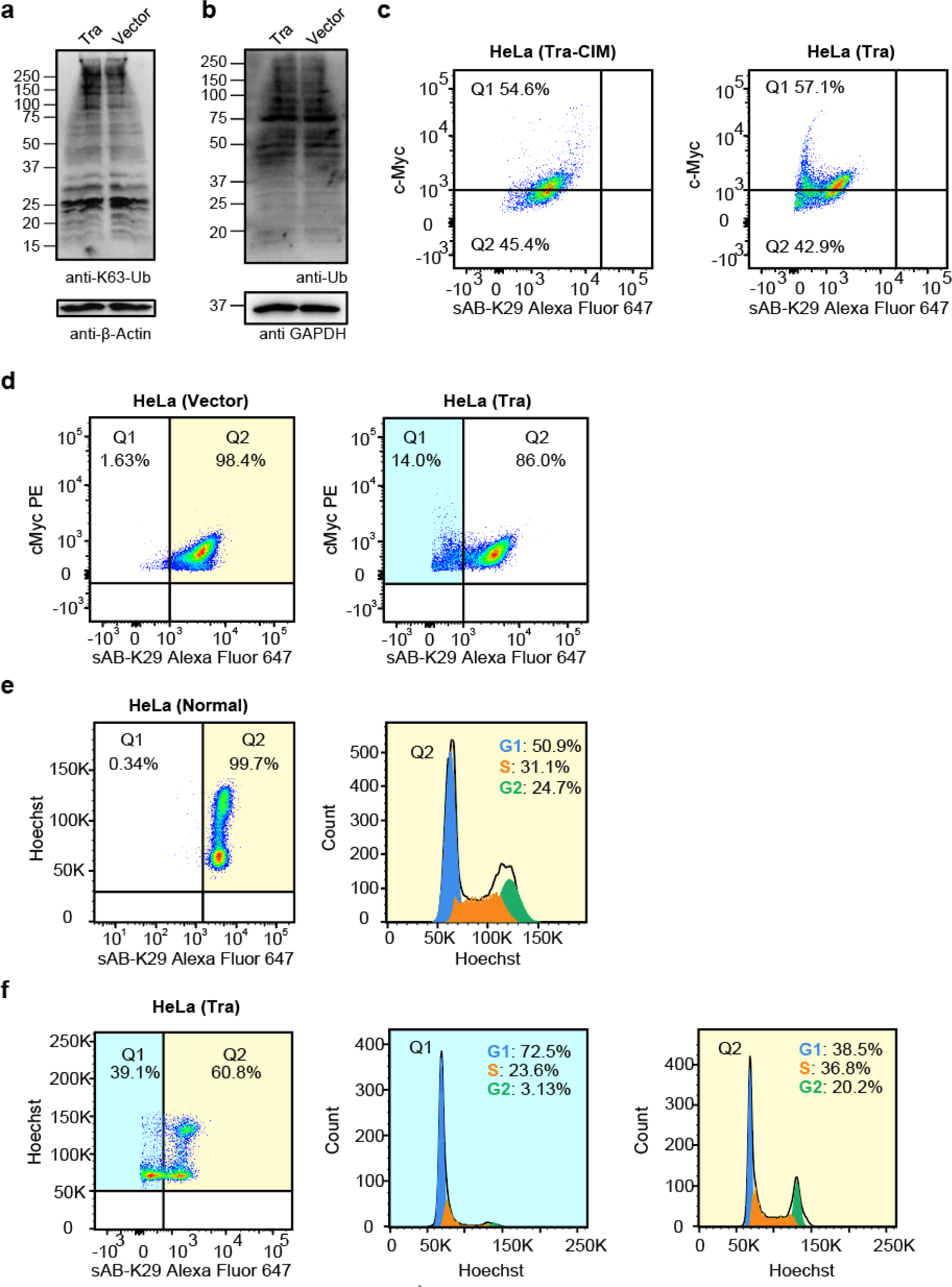
Reduced K29-linked ubiquitination led to G1/S arrest. **a-b**, Western blots of K63-linked ubiquitin and total ubiquitin in HeLa cells transfected with the Tra construct or the empty vector. **c**, Cell cycle analysis of HeLa cells transfected with either the Tra or the Tra-CIM construct by flow cytometry. cMyc and K29-linked polyubiquitin channels are shown. Representative flow cytometry graphs from three biological replicates are shown. **c**, Cell cycle analysis of HeLa cells transfected with either the Tra construct or the empty vector by flow cytometry. cMyc and K29-linked polyubiquitin channels are shown. Note that the cMyc tag was also expressed in cells transfected with the empty vector. Representative flow cytometry graphs from two biological replicates are shown. **e**, Cell cycle analysis of normal HeLa cells by flow cytometry. **f**, Cell cycle analysis of Hela cells transfected Tra construct for 27 h using flow cytometry. Representative flow cytometry graphs from three biological replicates are shown (a total of 5 injections). Highlighted areas were subjected to cell cycle analysis.

**Extended Data Fig. 7.**
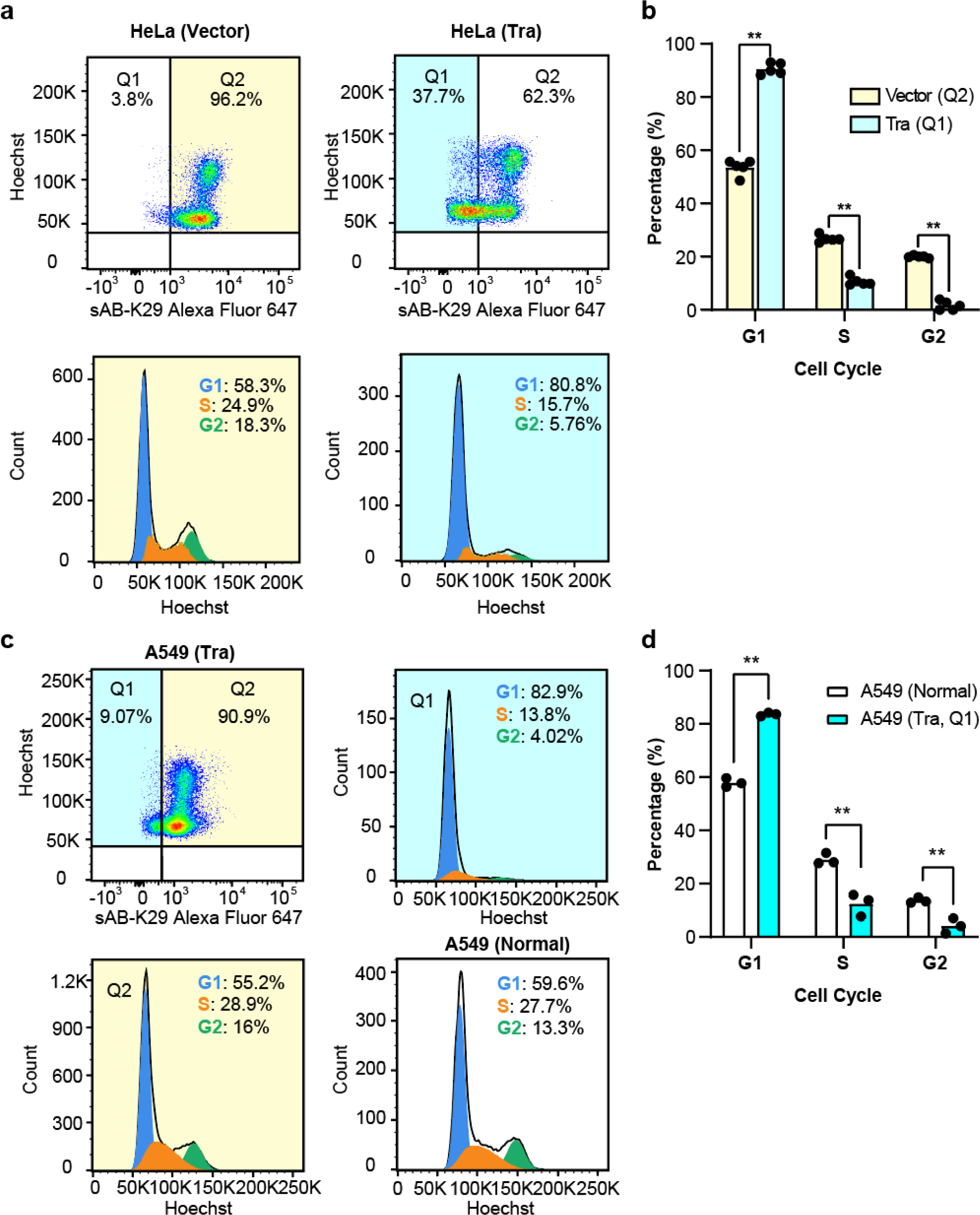
Quantitative analysis of reduction in K29-linked ubiquitination leading to G1/S arrest. **a**, Cell cycle analysis of HeLa cells transfected with Tra or an empty vector by flow cytometry. Representative flow cytometry graphs from two biological replicates are shown (a total of 5 injections). Highlighted areas were subjected to cell cycle analysis. **b**, Quantification of the cell cycle analysis results in panel **a**. Two biological replicates (a total of 5 injections) were included in this experiment. (**, p < 0.01 by two-tailed Student’s *t*-test). Error bars represent standard deviations. **c**, Cell cycle analysis of A549 cells transfected with either the Tra construct or empty vector by flow cytometry. **d**, Quantification of the A549 cell cycle analysis results in panel C. Error bars represent standard deviations from three injections (**, p < 0.01 by two-tailed Student’s *t*-test).

**Extended Data Fig. 8.**
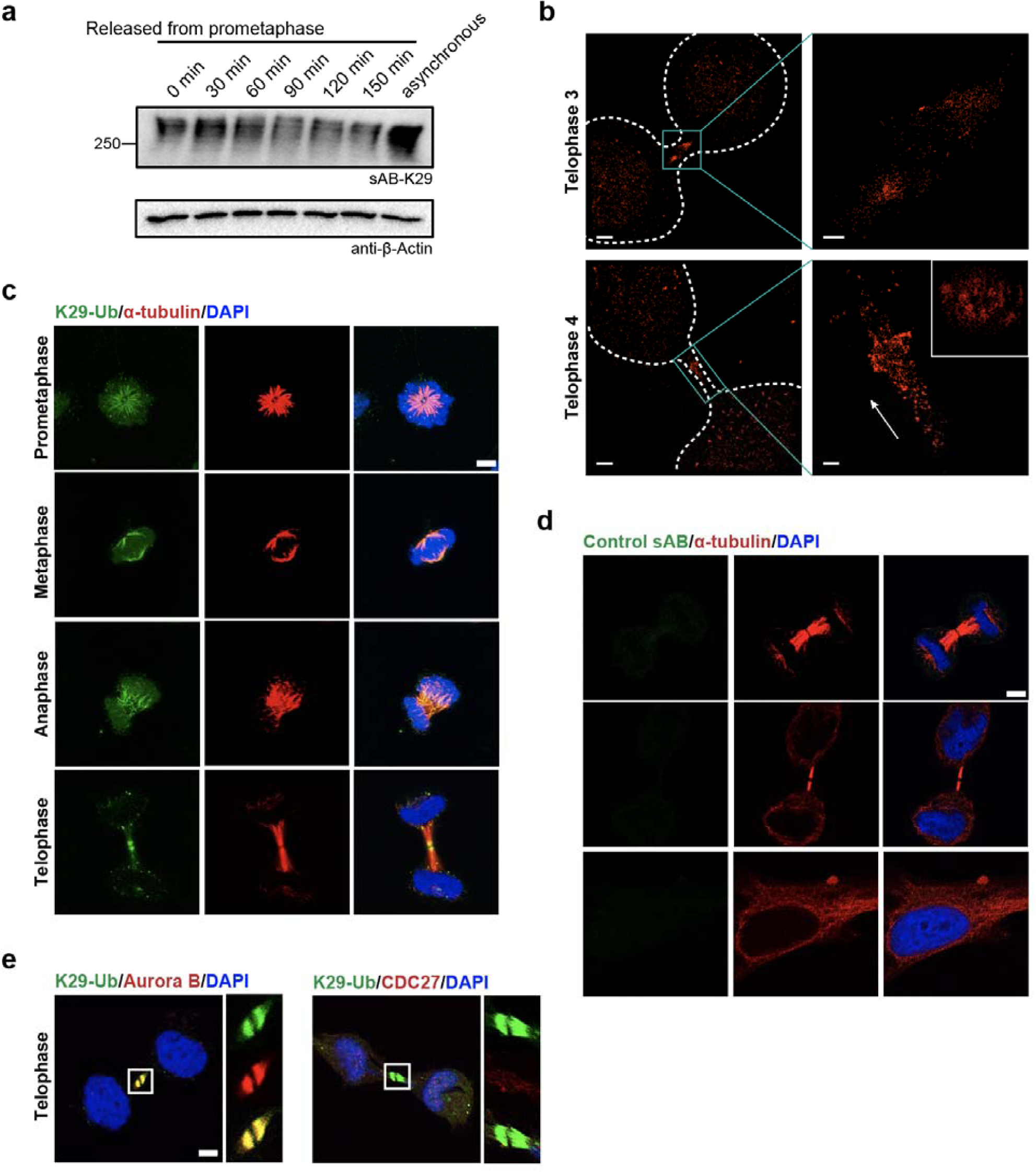
K29-linked ubiquitination is involved in mitosis. **a**, Western blot analysis of K29-linked ubiquitination during mitosis. HeLa cells were synchronized to prometaphase by treatment with thymidine and nocodazole. Aliquots of cells were taken and lysed at the indicated time points, followed by Western blotting using sAB-K29 as the primary binder. **b**, 3D STORM images of K29-linked polyubiquitin at telophase 3 and 4. The orthographic projection image of the highlighted area in telophase 4 (bottom cyan box) along the long axis (white arrow) is shown in the upper right inset. Rendering was conducted by Visual Molecular Dynamics (VMD) with GLSL rendering, Orthographic display, and Surf drawing. Movies showing rotation are presented as Movies 1A and 1B. Scale bars in the left images correspond to 2 μm. Scale bars in the right images correspond to 500 nm. **c**, Immunofluorescent staining of HeLa cells at different stages of mitosis. Costaining for K29-linked ubiquitin with α-tubulin is shown. Cells were pre-extracted before fixation. **d**, Immunofluorescent staining of HeLa cells with a control sAB (against maltose-binding protein). Costaining for α-tubulin is shown. Cells were pre-extracted before fixation. **e**, Immunofluorescent staining of HeLa cells at the telophase of mitosis. Costaining for K29-linked ubiquitin with either Aurora B or CDC27 (a negative control) is shown. Cells were pre-extracted before fixation. Scale bars in panels **c**, **d**, and **e** correspond to 5 μm.

**EXTENDED DATA Fig. 9.**
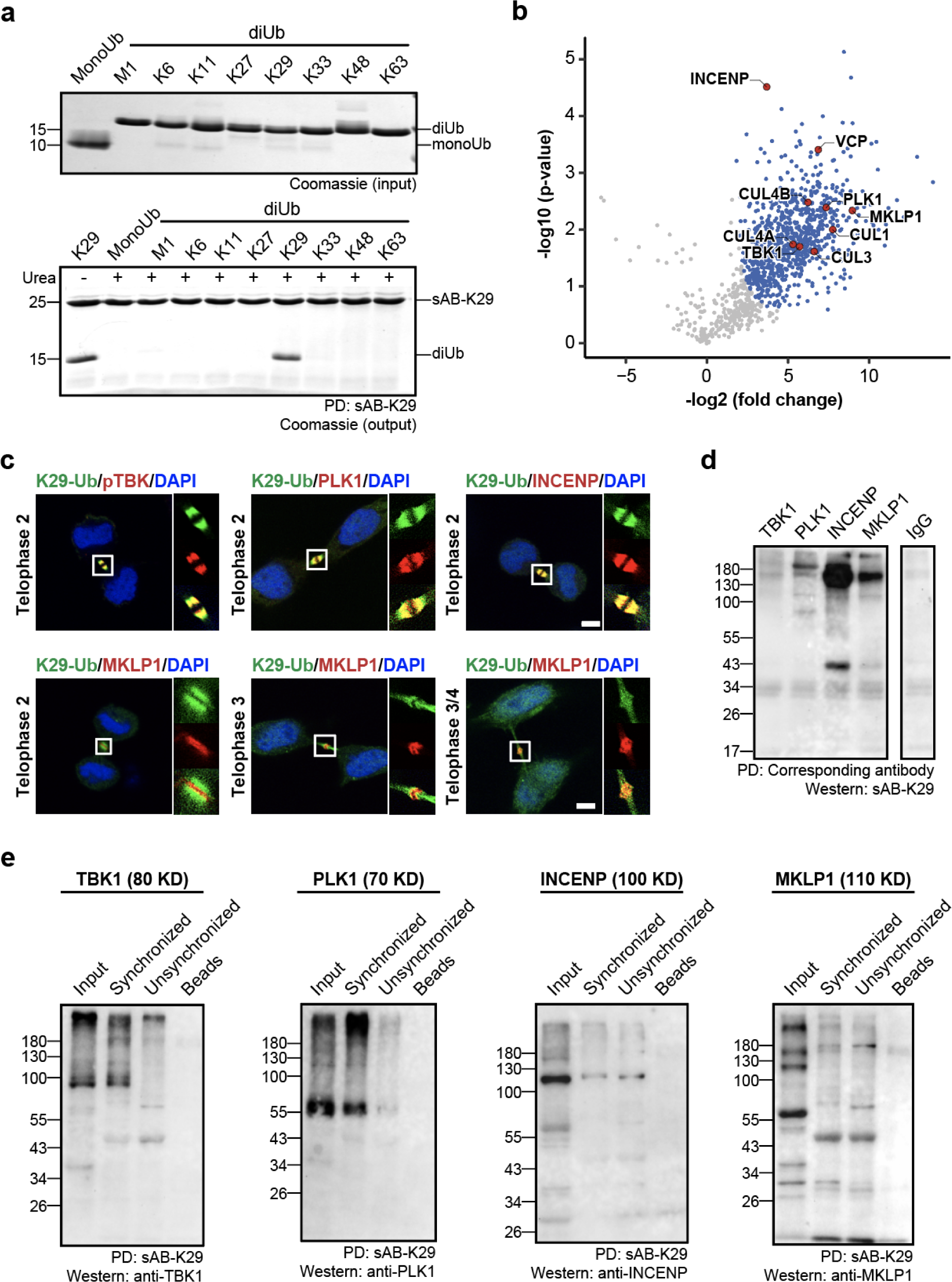
Midbody proteins involved in K29-linked polyubiquitination. **a**, sAB-K29 could be used to specifically pull down K29-linked diUb under denaturing conditions (2 M urea). **b**, Volcano plot of the quantitative mass spectrometry results after pull-down experiments in HeLa cells synchronized to the telophase of mitosis using sAB-K29 under denaturing conditions (1 M urea). Significant hits (FDR < 0.05) are colored blue, with those involved in midbody assembly highlighted in red. **c**, Immunofluorescent staining of HeLa cells at the telophase of mitosis. Costaining for K29-Ub with pTBK, PLK1, INCENP, and MKLP1 is shown. Scale bars correspond to 5 μm. **d**, Immunoprecipitation of synchronized HeLa cells under denaturing conditions using antibodies against TBK1, PLK1, INCENP, and MKLP1. Western blotting was performed using sAB-K29 as the primary binder. Anti-IgG was used in the control experiment. **e**, Immunoprecipitation of HeLa cells (either synchronized to the telophase of mitosis or unsynchronized) using sAB-K29. Antibodies against TBK1, PLK1, INCENP, and MKLP1 were used for Western blotting.

**Extended Data Fig. 10.**
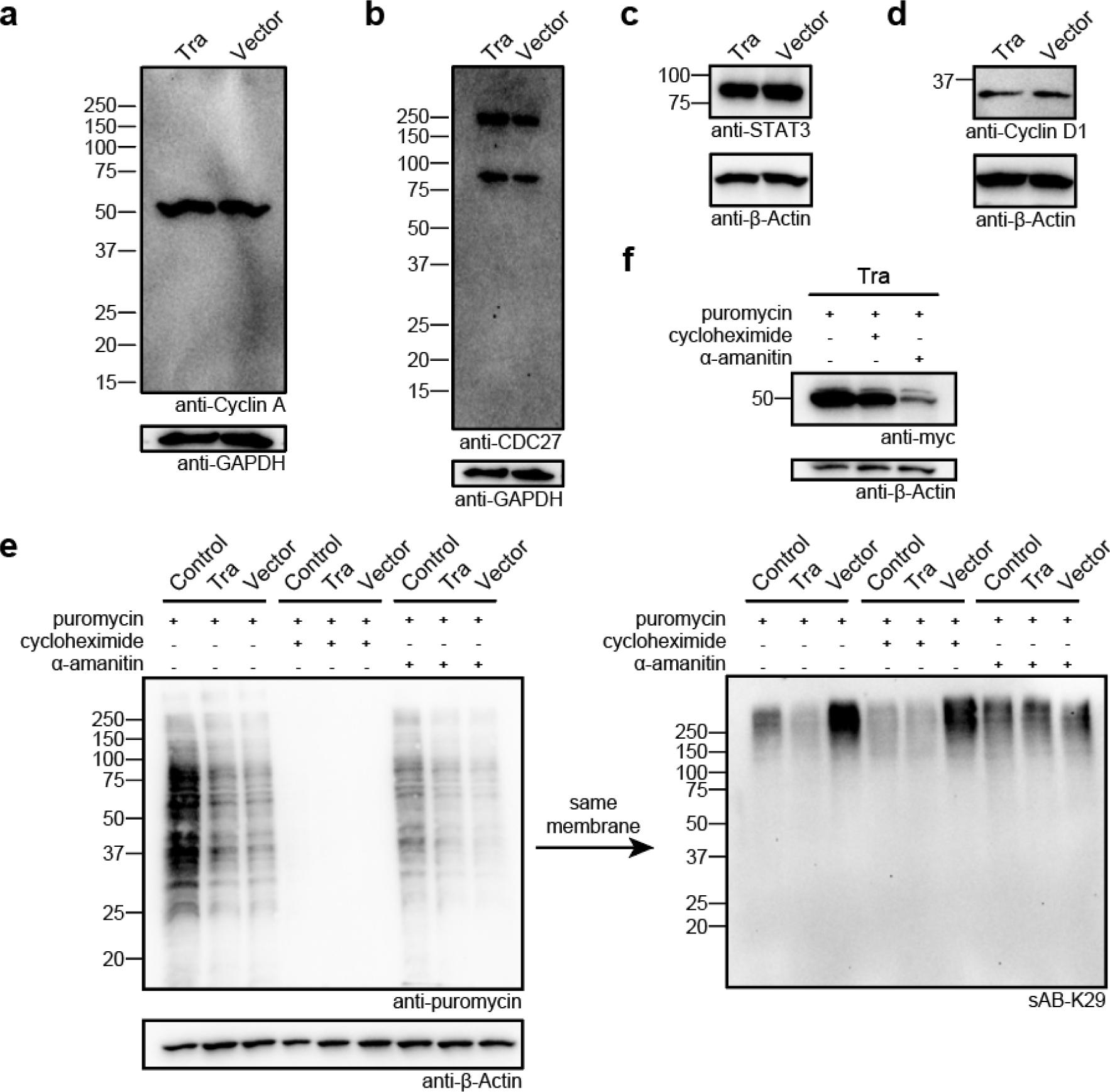
Western blot analysis of HeLa cells after the level of K29-linked ubiquitination was reduced using the Tra construct. **a-d**, Western blots of CyclinA, CDC 27, STAT3, and Cyclin D1 in HeLa cells transfected with the Tra construct or the empty vector. **e**, New protein synthesis was not affected after K29-linked ubiquitination was reduced in HeLa cells. HeLa cells were transfected with the Tra construct or control vector and treated with puromycin, cycloheximide, or α-amanitin. Western blotting was performed using anti-puromycin antibody or sAB-K29. **f**, Expression level of the Tra construct in HeLa cells treated with puromycin, cycloheximide, or α-amanitin. Western blotting was performed using an anti-cMyc antibody.

---

**Supplementary Movie 1**

3D STORM visualization (rotation by a shorter axis) of K29-linked ubiquitin in the midbody of dividing HeLa cells. Rendering was conducted by Visual Molecular Dynamics (VMD) with GLSL rendering, Orthographic display, and Surf drawing.

**Supplementary Movie 2**

3D STORM visualization (rotation by a longer axis) of K29-linked ubiquitin in the midbody of dividing HeLa cells. Rendering was conducted by Visual Molecular Dynamics (VMD) with GLSL rendering, Orthographic display, and Surf drawing.

